# Implication of novel variants of *BMP2* in isolated congenital heart disease: Functional characterization by *in silico* and *invitro* approaches

**DOI:** 10.1101/2025.08.29.672127

**Authors:** Jyoti Maddhesiya, Ritu Dixit, Ashok Kumar, Bhagyalaxmi Mohapatra

## Abstract

Bone morphogenic protein2 (BMP2), a member of TGF-β super-family, known to play a wide range of roles during embryonic development, particularly in the formation of bone/skeleton, differentiation of neurons, skeletal muscle, and development of cardiac valve septa and outflow tract. *BMP2* haploinsufficiency is reported to cause multiple congenital malformations including cardiac defects mainly endocardial cushion formation and chamber specification. To investigate the functional relevance of *BMP2* variations in isolated CHD cases, we performed genetic screening of *BMP2* in 285 CHD probands along with 400 healthy controls by Sanger’s method. Five non-synonymous variants namely, an already known variant p.Ser37Ala in N-terminal region, one nonsense variant, p.Lys241X in pro-peptide region and three missense variants p.His321Leu, p.Glu328Lys and p.Ser351Cys in mature domain, were identified in 8 unrelated CHD cases. *In vitro* functional analysis by western blotting depicted an increase in phosphorylation of SMAD1/5 due to all five variants. Furthermore, overexpression of cardiac-specific downstream target genes namely *Smad1, Smad4, Smad5, Nkx2*.*5, Gata4* and *Irx4* of the BMP pathway was observed in response to all the variants. Luciferase assay also validated the enhanced expression of multiple downstream promoters *Id1-luc, Id3-luc, Tlx2-luc, and p(SBE)*_*4*_*-luc*. Additionally, computational analysis of RNA structural features and protein secondary and tertiary structural changes also highlighted the increased activity of mutants, possibly due to enhanced interactions of mutant proteins with their binding partners owing to more stable structures. Overall, this is the first study which characterized the functional association of *BMP2* variants with the pathogenesis of CHD by *in vitro* and *in silico* methods.

## 1. Introduction

Congenital heart disease (CHD) is the most common inborn heart defect with a high rate of morbidity and mortality and is estimated to affect nearly 1% of newborns. Over the past few decades, more than 100 genes have been associated with CHD. Mutations in these genes prompt the identification of molecular markers and reinforce our understanding of cardiac development.

Heart development is a complex biological process that depends on spatiotemporal interactions between different multipotent transcription factors. The expression of such factors, namely *NKX2*.*5, GATA4/5/6*, and *TBX20/5/1*, leads to mutual interactions that results in the proper development of fully functional four-chambered heart. Involvement of synchronized signaling networks which incorporate transforming growth factor-β (TGF-β), bone morphogenetic protein (BMP), Wnt family member (WNT), nodal growth differentiation factor (NODAL), and fibroblast growth factor (FGF) mutually co-ordinate during cardiogenesis. Of these, the BMP subclass of proteins that form either homo-or heterodimeric complexes to transduce signals, and have been acclaimed for their significant role in embryogenesis, dorsoventral axis formation (Panopoulou et al. 1998) as well as growth, differentiation, proliferation and migration of different types of cells of mesenchymal origin that induce cranio-skeletal, neuronal and cardiac development. The direct role of Bmps in the establishment of the heart tube and looping has been demonstrated in zebrafish (Chen et al. 1997). Similarly, Bmps are broadly expressed in the anterior endoderm and cardiac mesoderm prior to cardiac differentiation during heart development in Xenopus (Hemmati□Brivanlou and Thomsen 1995; Wang et al. 1997). The involvement of BMP2 and BMP4 via HAND1 mediated outflow tract formation has been reported in a mouse model (Zheng et al. 2021).

Bone morphogenic protein2 (BMP2), commonly known for its osteogenic function, also plays a critical role in embryonic mesoderm formation and heart development. Zhang and Bradley (1996) have shown that the BMP2 knock out (KO) mice are embryonic lethal showing malformation of heart along with abnormality in the development of amnion and chorion (Zhang and Bradley 1996). BMP2 expression was detected in the extraembryonic mesoderm as well as in the myocardium. Multiple studies have affirmed the critical role of BMP2 during epithelial-mesenchymal transformation (EMT) in endocardial cushion formation, that gives rise to the heart valve and septa (Sugi et al. 2004; Rivera-Feliciano and Tabin 2006). Conditional KO of BMP2 in cardiac progenitors prevents EMT, and in the myocardium of the AVC shows defects in formation of cardiac jelly and defective AV cushion morphogenesis are observed (Ma et al. 2005). BMP2 in the myocardium activates hyaluronic acid synthetase 2 (HAS2) to produce the cushion extracellular matrix required for EMT. BMP2 has also been shown to induce the expression of early cardiac key markers NKX2.5 and GATA4 within anterior mesodermal cells in proximity to the heart-forming region (Schultheiss et al. 1997; Ladd et al. 1998; Prall et al. 2007; Si et al. 2014).

The association of *BMP2* haploinsufficiency with human diseases was first reported in patients with 20p12.3 microdeletions exhibiting variable phenotypes including cleft palate, microcephaly, facial dysmorphism and hearing defects (Sahoo et al. 2011). Similar monoallelic pathogenic variants have also been reported in multiple patients in two independent studies showing craniofacial and skeletal dysmorphism, developmental delay and cardiac defects (Tan et al. 2017; Priestley et al. 2023).

In the present study, we examined a cohort of 285 isolated CHD patients and 400 healthy control individuals from the Indian population and identified five nonsynonymous heterozygous rare single nucleotide variants (SNVs). *In vitro* functional studies supported by *in silico* analysis demonstrated impaired BMP2 function.

## 2. Materials and Methods

### 2.1 Enrolment and clinical assessment of participants

The study cohort recruited for this study included 285 cases (111 females and 174 males) of isolated CHD probands (with a median age of 3 years, ranging from 0-30 years of age) and 400 ethnicity age-matched healthy controls (median age 3.7 years, ranging from 1 to 32 years of age) from the Department of Cardiology and the Department of Paediatric Medicine, SS Hospital, Banaras Hindu University (BHU), Varanasi, India. All subjects were thoroughly examined by chest X-ray, 2D-colour doppler echocardiography and ECG to diagnose different CHD phenotypes. The clinical phenotypes of the study subjects are tabulated in Supplementary Table 1. However, CHD patients with associated extracardiac anomalies and/or chromosomal abnormalities were excluded from the study. The ‘Institutional Human Ethical Committee’ of the University approved the study design.

### 2.2 Genetic screening and analysis of identified variants

For mutational screening, peripheral venous blood (3-5 ml) from patients, their available relatives as well as healthy age-matched control individuals after obtaining informed consent. Genomic DNA was isolated from leukocytes of collected blood samples according to standard ethanol precipitation method. Protein-coding regions as well as the flanking exon-intron boundaries of *BMP2* (NG_023233.1), were amplified by polymerase chain reaction (PCR) for all CHD cases and controls. The amplified products were enzymatically purified by Exonuclease I (USB Products, Affymetrix, Inc. USA) and recombinant Shrimp Alkaline Phosphatase (USB Products, Affymetrix, Inc.). The purified products were bi-directionally sequenced by Sanger’s method using Big Dye® Terminator v3.1 Sequencing Kit (Applied Biosystems, Inc. USA) using an ABI PRISM 3130 genetic analyzer (Applied Biosystems, USA) according to manufacturer’s protocol. Finch TV software (http://www.geospiza.com/ftvdlinfo.html, Geospiza) was used to analyze the DNA sequence chromatogram. The identified variants were confirmed by re-sequencing the same samples with reverse primers as well as by sequencing freshly amplified PCR products from the DNA of the respective subjects. The inheritance pattern was also examined by parental genotyping. Sequence novelty of identified variants was checked by exploring ClinVar (http://www.ncbi.nlm.nih.gov/clinvar/) database, gnomAD (https://gnomad.broadinstitute.org/), Indigenome (https://clingen.igib.res.in/indigen/), 1000Genome (https://www.internationalgenome.org/) and GenomeAsia100K (https://browser.genomeasia100k.org/) browser.

### 2.3 In silico analysis

To study the effects of variants on the stability, potential pathogenicity, structure, and functions of wild-type (WT) and mutant proteins (muteins), different online *in silico* tools were searched. The reference mRNA sequence (NM_001200.4) and protein sequence (NP_001191.1) from NCBI (https://www.ncbi.nlm.nih.gov/genbank/) and Protein database (https://www.ncbi.nlm.nih.gov/protein/) were retrieved to perform all the *in silico* analyses.

#### 2.3.1 Evolutionary conserved amino acids

By retrieving the reference protein sequence from the Protein database (https://www.ncbi.nlm.nih.gov/protein/), BMP2 protein (NP_001191.1) was aligned across different orthologs of mammalian and non-mammalian species including *H*.*sapiens, P*.*troglodytes, M*.*mulatta, C*.*lupus, B*.*taurus, M*.*musculus, R*.*norvegicus, G*.*gallus, D*.*rerio* to check the evolutionary conservation of substituted amino acids by ‘Homologene’ tool of NCBI (http://www.ncbi.nlm.nih.gov/homologene).

#### 2.3.2 Predicting the disease-causing potential of BMP2 variants

The potential pathogenicity of non-synonymous variations that alter the structure, stability and function of proteins was predicted using an integrated database Varcard2 (http://www.genemed.tech/varcards2/#/index/home) which encompasses more than 10 bioinformatics tools such as SIFT, Polyphen2_HVAR, LRT, MutationTaster, PROVEAN, VEST3, M_CAP, CADD, DANN, GenoCanyon, ClinPred, ReVe and REVEL etc. The predictions of these *in silico* tools are based on different algorithms and parameters such as the nature of R group, the conservation of nucleotides and amino acid residues.

#### 2.3.3 Alteration in physicochemical properties of BMP2 muteins

The physicochemical properties of proteins determine their tertiary structure and function. The predictions of changes in different physicochemical properties (alpha-helix, hydrophobicity, refractivity, and relative mutability) were checked by Protscale (https://web.expasy.org/protscale/), a bioinformatic server at Expasy. To check the effect of variants on the above-listed physicochemical properties, a comparative analysis was performed for WT versus muteins of BMP2.

#### 2.3.4 Prediction of mutational effect on RNA features

To further delve into the mutational analysis of the structural alterations induced by silent mutations, the MutaRNA tool (version (5.0.10) was utilized (Miladi et al. 2020). This analysis encompassed assessing the intramolecular base pairing potential, base pairing probabilities of the mutant (MUT) mRNA, and assessment of accessibility (single-strandedness) in comparison to the WT counterpart (Bernhart et al. 2011). By amalgamating remuRNA (Salari et al. 2013) and RNAsnp, the changes induced by mutations in RNA structures can be statistically figured out.

#### 2.3.5 Conformational change in the 2-D and 3-D structures of muteins

Alterations in the secondary structures of proteins introduced by non-synonymous variations were predicted using online server Psipred (http://bioinf.cs.ucl.ac.uk/psipred/). For further validating these conformational changes, modelling of tertiary structures of BMP2-WT and all muteins were performed with a confidence score of more than 90% by Alphafold2 googlecolab, (https://www.nature.com/articles/s41586-021-03819-2). PyMOL Molecular Graphics System, Version 3.1 (Schrödinger, LLC.) was used to superimpose the modelled WT and muteins structures and visualize the differences in the conformation of the superimposed structures. The visualized differences were further validated by the global RMSD value which was calculated by UCSF ChimeraX.

### 2.4 Characterizing BMP2 variants by in vitro analysis

#### 2.4.1 Cloning of BMP2 and site-directed mutagenesis

Full-length *Homo sapiens* cDNA of *BMP2* was purchased from “Addgene” (Cat No. #137909, Watertown, MA, USA). The coding region was PCR-amplified using PfuUltra high-fidelity DNA polymerase (Stratagene, Santa Clara, CA, USA) and subcloned into the pcDNAv3.1/NT-GFP TOPO TA vector (Invitrogen, Carlsbad, CA, USA). To prepare the mutant constructs, the identified mutant nucleotides (c.109T>G, c.721A>T, c.962A>T, c.982G>A, and c.1052C>G) were introduced into the WT-BMP2-pcDNAv3.1/NT-GFP (BMP2_WT_GFP) by using Quick change II XL site directed mutagenesis kit (Agilent Technologies Inc.) with a complementary pair of designed mega primers. These mutant constructs were transformed into TOP10 competent cells followed by plasmid isolation (Qiagen midi kit, Germany). The fidelity of all mutant constructs (*BMP2*_Ser37Ala_GFP, *BMP2*_Lys241X_GFP, *BMP2*_His321Leu_GFP, *BMP2*_Glu328Lys_GFP, *BMP2*_Ser351Cys_GFP) was verified, by Sanger sequencing.

#### 2.4.2 Cell culture and transfection

*In vitro* functional analysis was performed using the mouse pluripotent embryonic carcinoma cell line P19 (kindly gifted from Dr. Ramkumar Sambasivan, InStem, India) and cardiomyoblast H9c2 (purchased from NCCS, Pune, India). P19 cell line was grown in α-MEM (Gibco, Life technologies Corp.) supplemented with 10% fetal bovine serum (HIMEDIA, India.) with 100 units/mL penicillin and 100 units/mL streptomycin and H9c2 (due to cardiomyoblast origin and better morphology, mainly used for immunostaining experiments) were cultured in DMEM supplemented with 10% serum at 37°C with 5% CO2 in a moist chamber. After 24 h of cell seeding, transfection was carried out at 30-40% confluency with FuGENE 6 transfection reagent (Promega Corp., IN, USA) as per the manufacturer’s instruction.

#### 2.4.3 Immunocytochemistry

Through immunocytochemistry, the cellular localization of BMP2 GFP was visualized in WT and muteins. The experiment was performed using H9c2 cell lines in a 6-well culture plate with each well containing glass cover slips. Transfection was performed with either *BMP2*_WT_GFP and mutant constructs (1 μg) 24 h after seeding. Cells were washed with chilled 1X PBS after 48 h of transfection followed by immediate fixation with 4% paraformaldehyde (PFA) for 15 min. The fixed cells were treated with 0.5% Triton-X and incubated for 30 min for permeabilization. Cells were washed with 1X PBST (5 times) followed by staining of nuclei with DAPI (Sigma-Aldrich) and the cytoskeleton with phalloidin (Sigma-Aldrich). Confocal imaging was carried out using Leica SP8 STED Laser Scanning Super Resolution Microscope System, Germany. Image analysis was performed using ImageJ software and assembled by Adobe Photoshop software.

#### 2.4.4 Western Blotting

The expression of BMP2 muteins was estimated by western blotting. P19 cells were grown in 6 well-culture plate followed by serum starvation for 12 h before transfection. Transfection was performed with 1 μg of BMP2 WT and mutant constructs followed by lysate preparation in RIPA Buffer 48 h post-transfection. Bradford assay was performed to quantify total proteins. Non-reducing samples (25 µg) were denatured for 5 min at 95°C in Laemmli’s loading buffer followed by resolving samples on 8% SDS polyacrylamide gel. Proteins were then transferred to a PVDF membrane (Bio-Rad Laboratories Inc, CA, US) and blocked with 5% non-fat dry milk in 1X TBST for 2 h at RT. Following blocking, the membrane was incubated with BMP2 specific primary antibody (R&D systems Inc., USA) for overnight at 4°C and the membrane was washed 8 times (5 min each wash) in 1X TBST. It was further incubated with horseradish peroxidase (HRP) conjugated goat anti-mouse IgG antibody (Genei, Germany) for 2 h at RT. After washing with 1X TBST five times (10 min each wash), the membrane was visualized using chemiDoc (Amersham^TM^ Imager 680, Japan) with an enhanced chemiluminescence detection kit (GE health Care, USA).

Further, the phosphorylation of SMAD1/5 (Ser463/465) was also assayed by transfecting cells with either *BMP2*_WT_GFP and mutant constructs (1 μg). After 48 h of transfection, whole cell lysates were prepared in RIPA buffer and quantified by Bradford method. The co-transfection experiment with ALK2 (1 μg) was also carried out with either *BMP2* WT or mutant constructs (1 μg each). Reduced samples (25 µg) were denatured for 5 min at 95°C followed by resolution on 8% SDS polyacrylamide gel. The protein samples were transferred to a PVDF membrane followed by blocking with 5% non-fat dry milk in 1X TBST for 2 h at RT. The blocked membrane was incubated with phospho-SMAD1/5 primary antibody (CST, USA) at 4°C overnight followed by washing with 1X TBST (3 times) and then incubated with HRP conjugated goat anti-rabbit IgG antibody (Genei, Germany) for 2 h at RT. The membrane was washed with 1X TBST (5 times) and chemiluminescence detection was performed using an ECL detection kit (GE health Care, US) using chemiDoc (Amersham^TM^ Imager 680, Japan).

#### 2.4.5 Dual luciferase reporter assay

To investigate the functional impact of BMP2 muteins using downstream target promoters such as *Id1-luc, Id3-luc, Tlx2-luc*, and *p(SBE)*_*4*_*-luc* as reporter genes, dual luciferase reporter assay was performed. P19 cells were grown in 24-well culture plates and transfection was performed after 24 h of cell seeding with 125 ng of *BMP2*_WT_GFP and mutant constructs along with 125 ng of each reporter plasmid and 25 ng Renilla luciferase vector as an internal control. For co-transfection experiments, 125 ng of ALK2 (one of the active receptors of BMP signaling) along with each reporter plasmid and *BMP2* constructs were co-transfected in independent sets of experiment. Cells were washed with chilled 1X PBS followed by lysate preparation in 1X passive lysis buffer (PLB) after 48 h of transfection. Luciferase activity was measured by Dual-Luciferase Reporter Assay System (Promega Corp, WI, USA) using Synergy/ HTX multi-mode reader (BioTek Instruments, Inc., USA). Three independent experiments were performed in triplicate for each sample for all the reporter plasmids and the pooled data plotted as mean fold change with standard error of mean.

Luciferase reporter plasmids used were kind gifts (*Id1-luc* from Maria Genander, Karolinska Institutet, Stockholm, Sweden; and *Id3-luc* from Daniel J. Bernard, Quebec, Canada; *Tlx2-luc* from Prof. Jeffrey L. Wrana, University of Toronto, Ontario, Canada; *p(SBE)*_*4*_*-luc* from Dr. Peter ten Dijke, Netherlands).

#### 2.4.6 Real-time PCR (qPCR) Assay

To perform real time PCR assay, P19 cells were grown in 6-well culture plate and transfected with *BMP2*_WT_GFP and mutant constructs (1 μg) at 50% confluency after 24 h. Total cellular RNA was extracted in TRI Reagent (Merck, Germany) after 48 h of transfection. RNA integrity was checked on 1% agarose gel followed by quantification using NanoDrop Micro UV/Vis spectrophotometer (ThermoFisher Scientific, USA). The RNA samples were treated with DNase I (ThermoFisher Scientific, USA) to remove any genomic DNA contamination. cDNA library was prepared from 2 µg of DNase I-digested RNA using Revertaid First Strand cDNA synthesis kit (ThermoFisher Scientific, USA) by following manufacturer’s protocol. The qPCR assay was performed to check the effect of *BMP2*_WT_GFP and mutant constructs on the expression level of downstream target genes *viz*., (*Smad1, Smad4, Smad5, Irx4, Nkx2*.*5, Gata4*) in real-time PCR machine (QuantStudio 5, Applied Biosystems, CA, USA) using SYBR green (Sigma-Aldrich, USA) as per manufacturer’s instruction. GAPDH was used as an internal control to normalized the data. Each independent experiments were performed in triplicate and the data were plotted as fold change with standard error of mean and the p value when <0.05 was considered statistically significant.

## 3. Results

### 3.1 Genetic variants of BMP2 and their genotype-phenotype correlation

The genetic evaluation of the coding exons and flanking intronic boundary regions of *BMP2*, in 285 sporadic CHD probands by Sanger sequencing, disclosed a total of 5 non-synonymous variants in 8 unrelated cases Fig. 1(b). Out of five variants, four were missense (c.109T>G; p.Ser37Ala, c.962A>T; p.His321Leu, c.982G>A; p.Glu328Lys and c.1052C>G; p.Ser351Cys) and one was nonsense (c.721A>T; p.Lys241X) variant. Four out of five variants were novel, not reported in any clinical database. These four novel variants were submitted in ClinVar database and their submission numbers are., p.Lys241X (SCV002568377), p.His321Leu (SCV002568378), p.Glu328Lys (SCV004022294) and p.Ser351Cys (SCV002568379). A previously reported missense variant (c.109T>G; p.Ser37Ala(rs2273073) was identified in the flanking region between the signal-peptide and pro-peptide domain, while c.962A>T; p.His321Leu, c.982G>A; p.Glu328Lys and c.1052C>G; p.Ser351Cys were identified in mature peptide domain. The nonsense variant c.721A>T; p.Lys241X was found towards the distal end of TGF-β pro-peptide domain. The variant p.Ser37Ala (rs2273073) was discovered in two unrelated male children, one of them diagnosed with complex CHD i.e., dTGA, and other one with ASD, VSD and PA. The nonsense variant p.Lys241X was identified in 2 years old female affected with PTA and VSD. The first mature peptide domain variant (p.His321Leu) was detected in a male child, with dTGA. Another mature peptide domain variant (p.Glu328Lys) was found in two unrelated male probands, one was 1.8 years, with VSD & PFO, and the other one was a 4 months old boy with large ASD & VSD along with biventricular enlargement. Genotype analysis of all four parental sample revealed all four carry homozygous WT allele (GG). The third mature peptide domain variant p.Ser351Cys was identified in two unrelated females. The first 12 years old female harbouring this variant was affected with ToF while the second 5 months old female diagnosed with VSD. None of the probands harbouring these *BMP2* variant had known family history of CHD or any other extracardiac abnormality. All the five identified variants were in heterozygous condition and none of the variant were detected in 400 ethnically aged-matched control (800 chromosomes). The coding positions along with amino acid change of these missense variants with their minor allele frequency (MAF) and clinical phenotypes are tabulated in Table 1. The heterozygous variants identified are shown in the electropherogram sequence along with the position on protein domain (Fig. 1a and b). Due to the low minor allele frequency of variants as well as significant association with CHD when compared to controls, these variants prompted us for further characterising their effect on structure and function of protein by *in silico* and *in vitro* approaches. The four novel non-synonymous variants (c.721A>T; p.Lys241X, c.962A>T; p.His321Leu, c.982G>A; p.Glu328Lys, c.1052C>G; p.Ser351Cys) were submitted to the ClinVar database (https://submit.ncbi.nlm.nih.gov/clinvar/) with received submission ID SUB11760578 and SUB13710701 (Table 1).

**Table 1.** Clinical phenotypes, position of variants in domain, ClinVar submission Ids and novelty.

| Nucleotide change | Amino Acid change | CHD Phenotype | Patient ID | Parental genotype | | MAF | | BMP2 Domain | ClinVar | gnomAD | Indigenome | 1000 Genome database | Genome Asia100K | I-mutant 2.0 ($\Delta\Delta G$, RI) |
| --- | --- | --- | --- | --- | --- | --- | --- | --- | --- | --- | --- | --- | --- | --- |
|  |  |  |  | Father | Mother | Cases (N=285) | Controls (N=400) |  |  |  |  |  |  |  |
| c.109T>G | p.Ser37Ala | dTGA, ASD, VSD, PA | CHD02 | NA | NA | 2 | 0 | Pro-peptide | Reported | rs2273073 | Reported | Reported | Reported | Decrease (0.01, 2) |
|  |  |  | CHD14 | NA | NA |  |  |  |  |  |  |  |  |  |
| c.721A>T | p.Lys241X | PDA | CHD433 | NA | NA | 1 | 0 | Pro-peptide | Novel (SCV002568377) | NR | NR | NR | NR | NA |
| c.962A>T | p.His321Leu | TGA | CHD186 | NA | AA | 1 | 0 | Mature domain | Novel (SCV002568378) | NR | NR | NR | NR | Increase (1.34, 3) |
| c.982G>A | p.Glu328Lys | VSD, VSD | CHD212 | GG | GG | 2 | 0 | Mature domain | Novel (SCV004022294) | NR | NR | NR | NR | Decrease (-0.22, 3) |
|  |  |  | CHD219 | GG | GG |  |  |  |  |  |  |  |  |  |
| c.1052C>G | p.Ser351Cys | ToF & ASD (PoF) | CHD13 | NA | NA | 2 | 0 | Mature domain | Novel (SCV002568379) | NR | NR | NR | NR | Decrease (-0.1, 5) |
|  |  |  | CHD24 | CC | CC |  |  |  |  |  |  |  |  |  |

**Abbreviations:** dTGA- dextro Transposition of great arteries, ASD- Atrial septal defects, VSD- Ventricular septal defects, PA- Pulmonary Atresia, PDA- Patent ductus arteriosus, ToF- Tetralogy of Fallot, PoF- Pentalogy of Fallot, MAF- Minor allele frequency, dbSNP- single nucleotide polymorphism database, NA- Not Available, NR- Not reported, RI- Reliability index

**Fig. 1.**
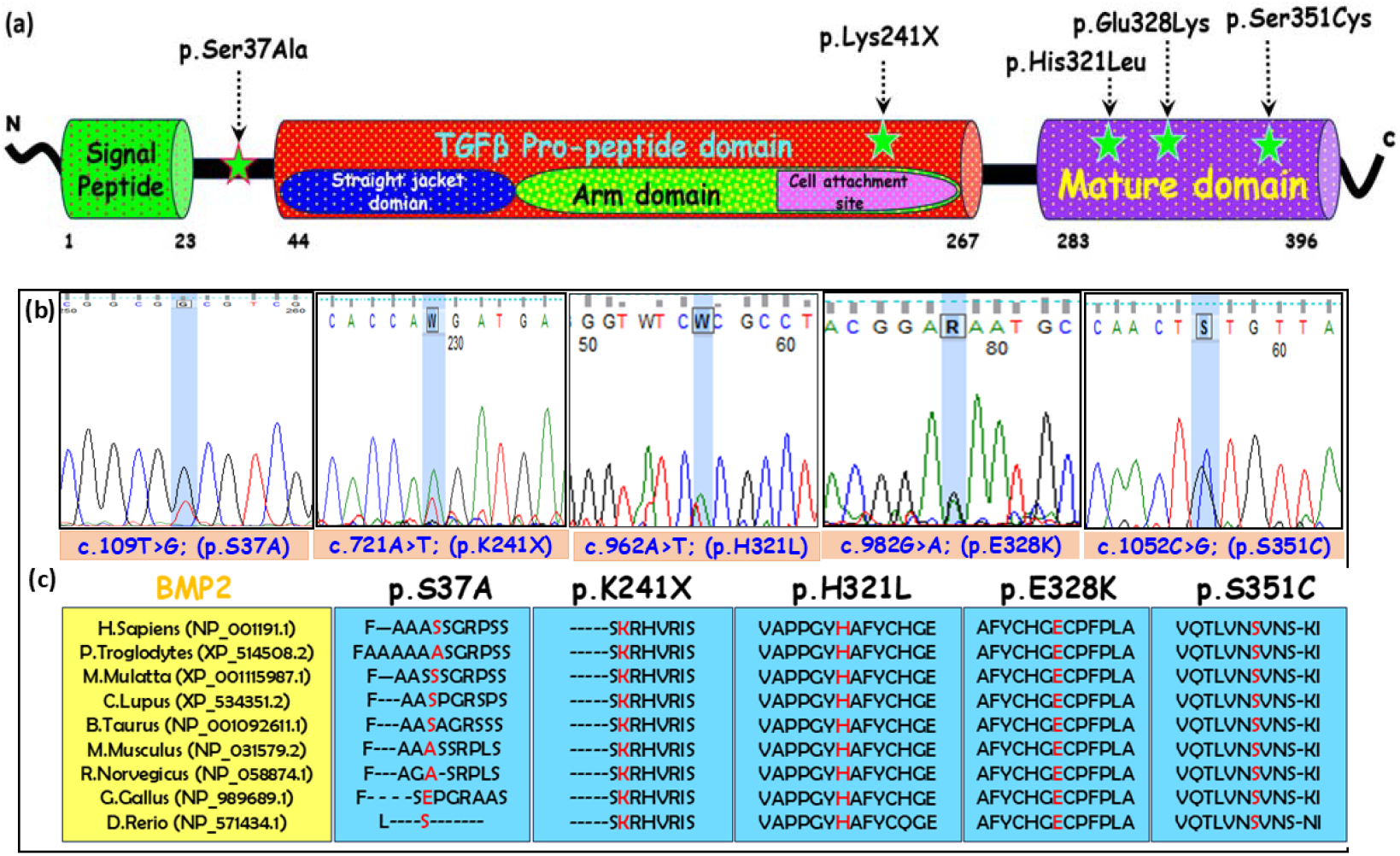
(1-a) Diagrammatic representation of BMP2 which is a 396 amino acids protein with different domains viz., signal peptide domain (1-23aa), TGFβ-pro-peptide domain (44-267aa), protease cleavage site (268-282aa) and mature peptide domain (283-396aa). All the missense identified variants are indicated with star in domain-wise location, (b) the chromatogram of sequence for the detected variants and (c) multiple sequence alignment of mutated amino acid (highlighted in red) across different species of vertebrate for all the five variants (p.Ser37Ala, p.Lys241X, p.His321Leu, p.Glu328Lys and p.Ser351Cys).

### 3.2 Analysis of phylogenetic conservation of BMP2 protein

To check evolutionary conservation of mutated amino acids for all the variants, multiple sequence alignment of BMP2 protein was performed across different species including *Homo sapiens* (NP_001191.1), *Pan troglodytes* (XP_514508.2), *Macca mulatta* (XP_001115987.1), *Canis lupus* (XP_534351.2), *Bos taurus* (NP_001092611.1), *Mus musculus* (NP_031579.2), *Rattus norvegicus* (NP_058874.1), *Gallus gallus*, (NP_989689.1), *Danio rerio* (NP_571434.1). The amino acid residue at position Lys-241, His-321, Glu-328 and Ser-351 showed 100 % conservation across above mentioned species. However, serine at position 37^th^ for the reported variant p.Ser37Ala is conserved only in few species. The conservation of substituted amino acid residues along with the flanking amino acid sequences showed in (Fig. 1c).

### 3.3 Disease-causing potential of BMP2 variants

Bioinformatic analysis were conducted to predict the disease-causing potential of the variants, by an online server VarCard2, that includes algorithm of more than 15 *in silico* tools. The prediction of most of the tools *viz*., fitCons, FATHMM_MKL, Eigen, VEST3, CADD and DANN for the pro-peptide domain variant p.Ser37Ala was tolerable. However, the other variants (p.Lys241X, p.His321Leu, p.Glu328Lys and p.Ser351Cys) were predicted to be pathogenic by ClinPred and damaging by fitCons, FATHMM_MKL, Eigen, VEST3, CADD and DANN. The predictions and the algorithmic scores by different *in silico* tools of all these variants were tabulated in Table 2.

**Table 2.** Predicted potential pathogenicity of the identified variants of BMP2 by different in-silico tools.

| Variant | ClinPred |  | fitCons |  | FATHMM_MKL |  | Mutation Taster |  | Eigen |  | VEST3 |  | CADD |  | DANN |  |
| --- | --- | --- | --- | --- | --- | --- | --- | --- | --- | --- | --- | --- | --- | --- | --- | --- |
|  | Prediction | Score | Prediction | Score | Prediction | Score | Prediction | Score | Prediction | Score | Prediction | Score | Prediction | Score | Prediction | Score |
| c.109T>G | benign | 0.00229<br>969 | Tolerable | 0.652 | Tolerable | 0.03 | Polymorphism | 1 | Tolerable | -2.008 | Tolerable | 0.055 | Tolerable | 0.001 | Tolerable | 0.351 |
| c.721A>T | NA | NA | Damaging | 0.707 | Damaging | 0.846 | Disease-causing | 1 | Damaging | 0.451 | NA | NA | Damaging | 37 | Damaging | 0.992 |
| c.962A>T | pathogenic | 0.97976<br>511 | Damaging | 0.707 | Damaging | 0.921 | Disease-causing | 1 | Damaging | 0.554 | Damaging | 0.716 | Damaging | 25 | Tolerable | 0.971 |
| c.982G>A | pathogenic | 0.94885<br>591 | Damaging | 0.707 | Damaging | 0.971 | Disease-causing | 1 | Damaging | 0.596 | Damaging | 0.66 | Damaging | 31 | Damaging | 0.999 |
| c.1052C>G | pathogenic | 0.99802<br>219 | Damaging | 0.707 | Damaging | 0.952 | Disease-causing | 1 | Damaging | 0.876 | Damaging | 0.798 | Damaging | 25.8 | Tolerable | 0.987 |

**Abbreviations:** ClinPred- ClinVar prediction, FATHMM MKL- Functional analysis through hidden Markov models, VEST3- Variant effect scoring tool, CADD- Combined annotation dependent depletion

### 3.4 Effect of variants on physicochemical properties of BMP2 protein

The different physicochemical properties such as alpha-helix, hydrophobicity, relative mutability, refractivity of proteins are greatly determined by the R-side chain of amino-acids. As the missense mutations change the amino-acid, the R-side chain of the newly introduced amino acid will be different from that of the replaced one. On the basis of these properties, we had performed *in silico* computational analysis by Protscale to compare these properties of mutants with the WT protein. We observed significant changes in alpha-helix due to variants p.Ser37Ala and p.Glu328Lys. The hydrophobicity was also significantly changed for all the muteins when compared with *BMP2* WT. Furthermore, other properties such as relative mutability and refractivity were also predicted and marked changes were observed in both the properties due to all the five variants (p.Ser37Ala, p.Lys241X, p.His321Leu, p.Glu328Lys and p.Ser351Cys) (Supplementary Fig. 1).

### 3.5 Effect of variants on RNA structural features

A remarkable change in the base-pairing probabilities of the MUT RNA, intra-molecular base pairing potential, and assessment of accessibility (single-strandedness) were observed in response to all the five variants Fig. 2. The circos plots were illustrating the base pairing probabilities with visualization of darker hue of grey depicting higher probability. Likewise, the dot plot matrices were portraying the base pair probability of *BMP2* WT and MUT RNA snippets with darker the dots specifying greater base pairing potential. The differential dot plot matrices were representing the change in base-pair probability {Pr (bp in WT) – Pr (bp in MUT)} which describe the difference in base-pairing patterns between WT and MUT RNAs at specific locations. A major modification in base-pairing probabilities was noted as displayed by the dot plots, circos plots and differential dot plots due to variants p.Ser37Ala and p.Ser351Cys. Nonetheless, remarkable alterations in base pairing probabilities exhibited by variants p.Lys241X, p.His321Leu and p.Glu328Lys as shown by the dot plots, circos plots and differential dot plots. Additionally, the accessibility profile of RNA structures (the probability of being unpaired at each base position which influences the RNA-protein or RNA-RNA interactions) have also depicted significant changes. Notably, variants p.Ser37Ala and p.Ser351Cys exhibits the most pronounced alteration in accessibility, following closely related variants p.Lys241X and p.His321Leu which show distinguished changes. Conversely, minor modifications caused due to variant p.Glu328Lys in RNA accessibility.

**Fig. 2.**
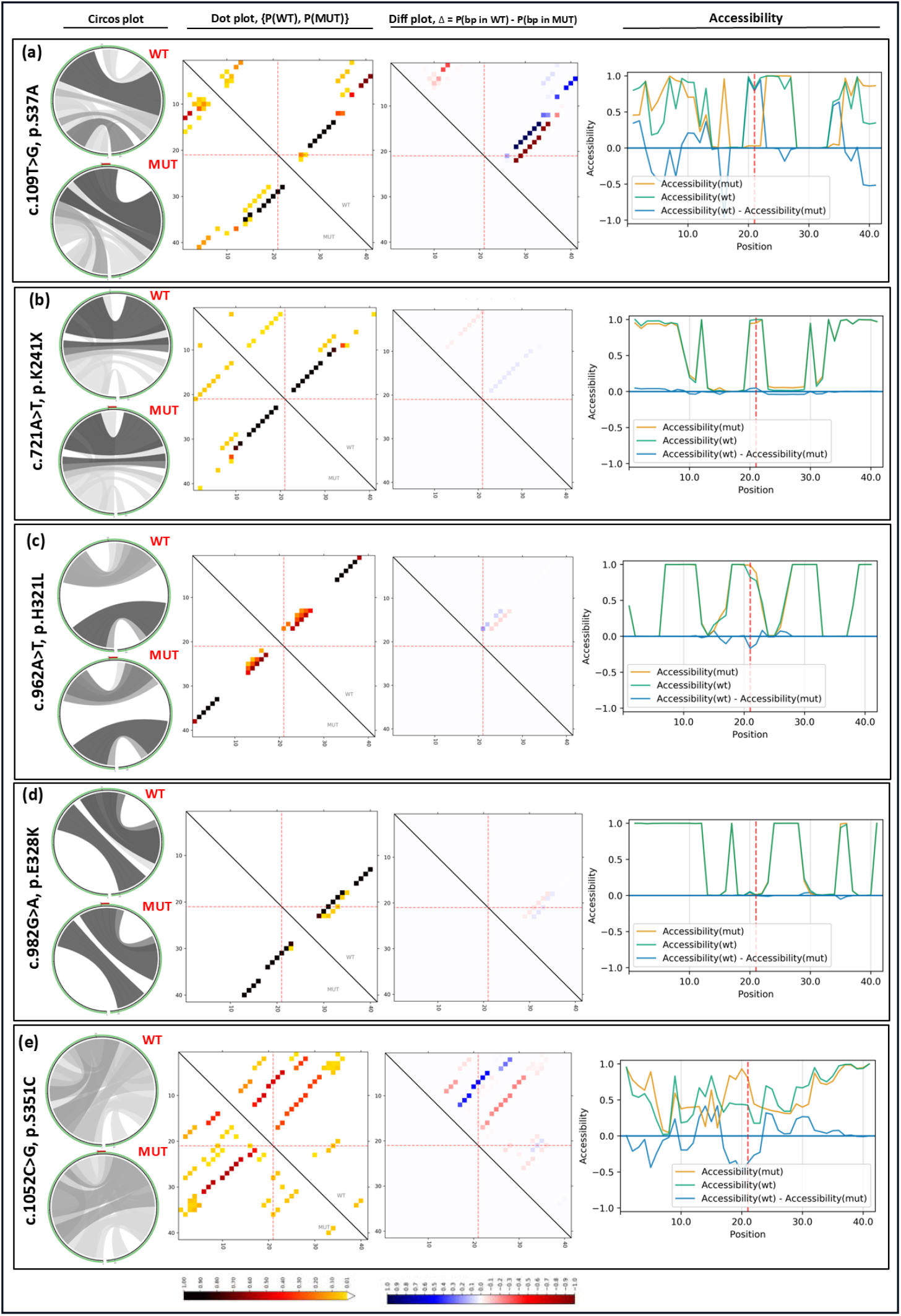
(a-e). Comparative analysis of RNA structural features of MUT with WT. The Circos plots represent the base – pairing probability of p(WT) and p(MUT) RNA structures, with WT plot at upper and MUT at lower. The sequence starting from 5’ end at the bottom-left and extending clockwise reaching to the 3’ end. The variant is at position 21 which is highlighted in red at the top of each MUT circos plot. The base-pairing of higher probability is illustrated in darker shades of grey. The dot plot is a heatmap-like representation of base-pair probability of p(WT) and p(MUT) structures. The base-pairing of higher probability is indicated in darker dots. The differential dot plot depicts the differences in base-pairing probabilities between the WT and MUT RNA [Pr (bp in WT) – Pr (bp in mut)]. The red color dots represent strong base-pairing while the blue color dots highlighting weak base pairing which results due to mutations. The last column graph demonstrating the accessibility profile in terms of unpaired probabilities of WT and MUT RNA. The blue line indicating the variation in accessibility (WT-MUT) with negative values depicting the Fig. 2(a-e). Comparative analysis of RNA structural features of MUT with WT. The Circos plots represent the base – pairing probability of p(WT) and p(MUT) RNA structures, with WT plot at upper and MUT at lower. The sequence starting from 5’ end at the bottom-left and extending clockwise reaching to the 3’ end. The variant is at position 21 which is highlighted in red at the top of each MUT circos plot. The base-pairing of higher probability is illustrated in darker shades of grey. The dot plot is a heatmap-like representation of base-pair probability of p(WT) and p(MUT) structures. The base-pairing of higher probability is indicated in darker dots. The differential dot plot depicts the differences in base-pairing probabilities between the WT and MUT RNA [Pr (bp in WT) – Pr (bp in mut)]. The red color dots represent strong base-pairing while the blue color dots highlighting weak base pairing which results due to mutations. The last column graph demonstrating the accessibility profile in terms of unpaired probabilities of WT and MUT RNA. The blue line indicating the variation in accessibility (WT-MUT) with negative values depicting the positions more prone to be unpaired in the MUT vs WT.

### 3.6 Effect of variants on secondary and tertiary structures of BMP2 proteins

Significant changes in the secondary structures of all BMP2 muteins were predicted (Supplementary Fig. 2). Comparing the predicted secondary structures of muteins due to variants p.Ser37Ala, p.Lys241X, p.His321Leu, p.Glu328Lys and p.Ser351Cys, significant changes were observed in both α-helix and β-sheet, as listed in Supplementary Table 2.

The tertiary structures of *BMP2* WT and muteins were modelled which revealed structural changes due to mutations as indicated by the aligned mutein structures versus WT. The remarkable changes in the structures of muteins were further supported by the global RMSD values. The reported variant with serine substituted by alanine illustrated major changes with supported RMSD value of 6.035 Å. The substitution of lysine by stop codon (p.Lys241X) showed that the mutated protein gets terminated after the incorporation of stop codon in the distal pro-peptide domain and the RMSD value was 10.116 Å. Furthermore, significant changes were also observed in the tertiary structures of the mature domain muteins viz., p.His321Leu, p.Glu328Lys, p.Ser351Cys when superimposed over WT with RMSD value of 6.293 Å, 6.216 Å, and 6.283 Å respectively indicating a greater conformational changes. These alterations in conformation of muteins implicate that mutated amino acid affect the tertiary structures of BMP2 protein. The aligned structures with WT protein in green and muteins in hotpink colours were presented in Fig. 3(a-e).

**Fig. 3.**
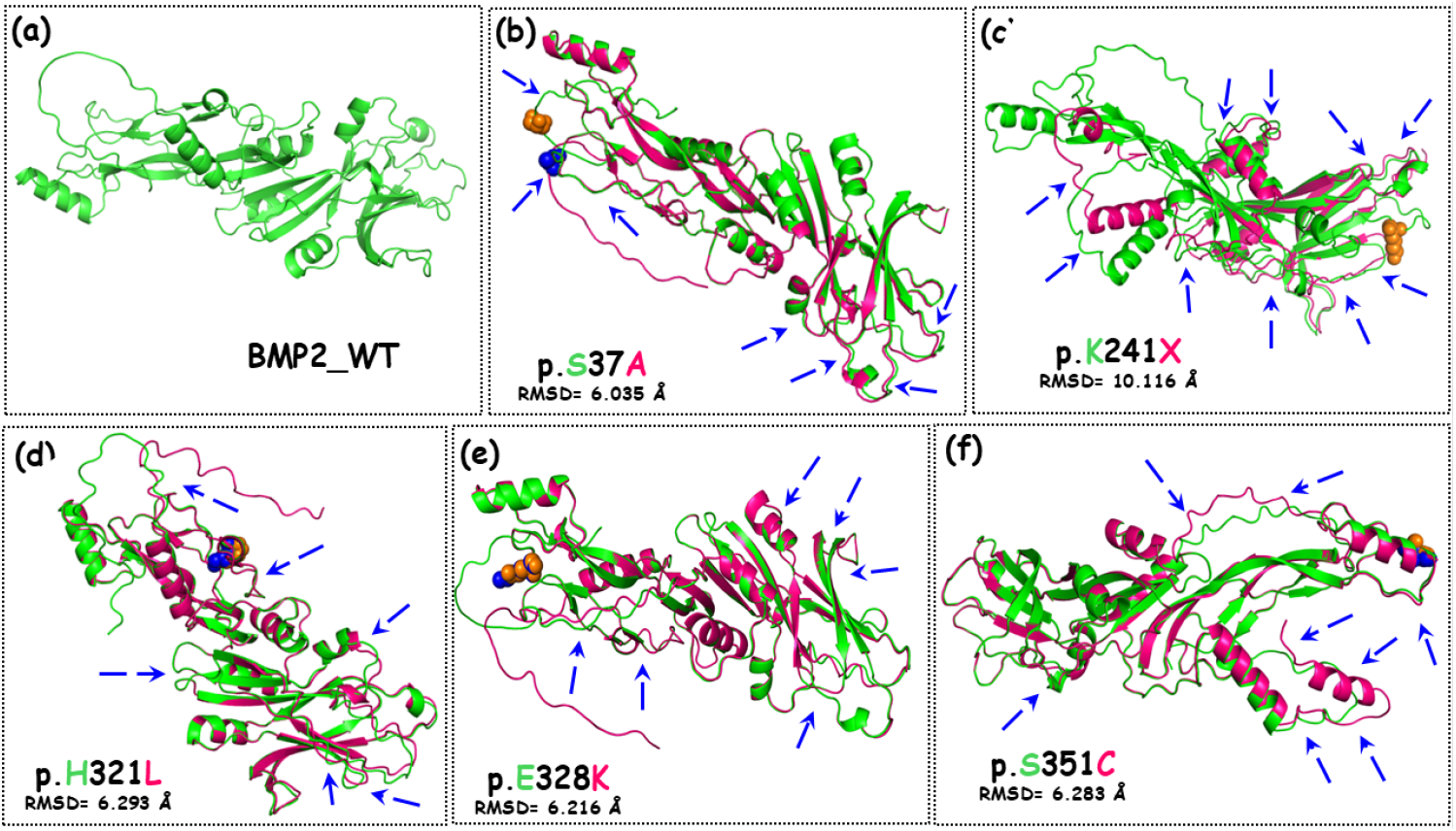
(a-e). Comparison based on the tertiary structures of *BMP2 WT* and muteins. Tertiary structural comparison of BMP2 muteins with WT, (a) modelled-tertiary structure of *BMP2 WT*, (b-e) superimposed tertiary structures of different muteins of BMP2 (p.Lys241X, p.His321Leu, p.Glu328Lys and p.Ser351Cys) over WT with supported global alpha carbon RMSD values. The tertiary structure of WT is shown in green colour while muteins are represented in hotpink with mutated residues in spheres. The conformational changes in the structures of muteins are indicated by arrows.

### 3.7 Impact of variants on the expression and cellular localization of BMP2 protein

The cellular localization of *BMP2 WT* and muteins was checked by immunofluorescence staining in H9c2 cell lines. The GFP-tagged *BMP2 WT* and muteins showed cytoplasmic as well as nuclear localization. Nonetheless, clustered accumulation of BMP2 in nucleus is intriguing. However, there was no significant change observed in the expression of muteins when compared to WT counterpart [Fig. 4(i-a1-f4)].

**Fig. 4.**
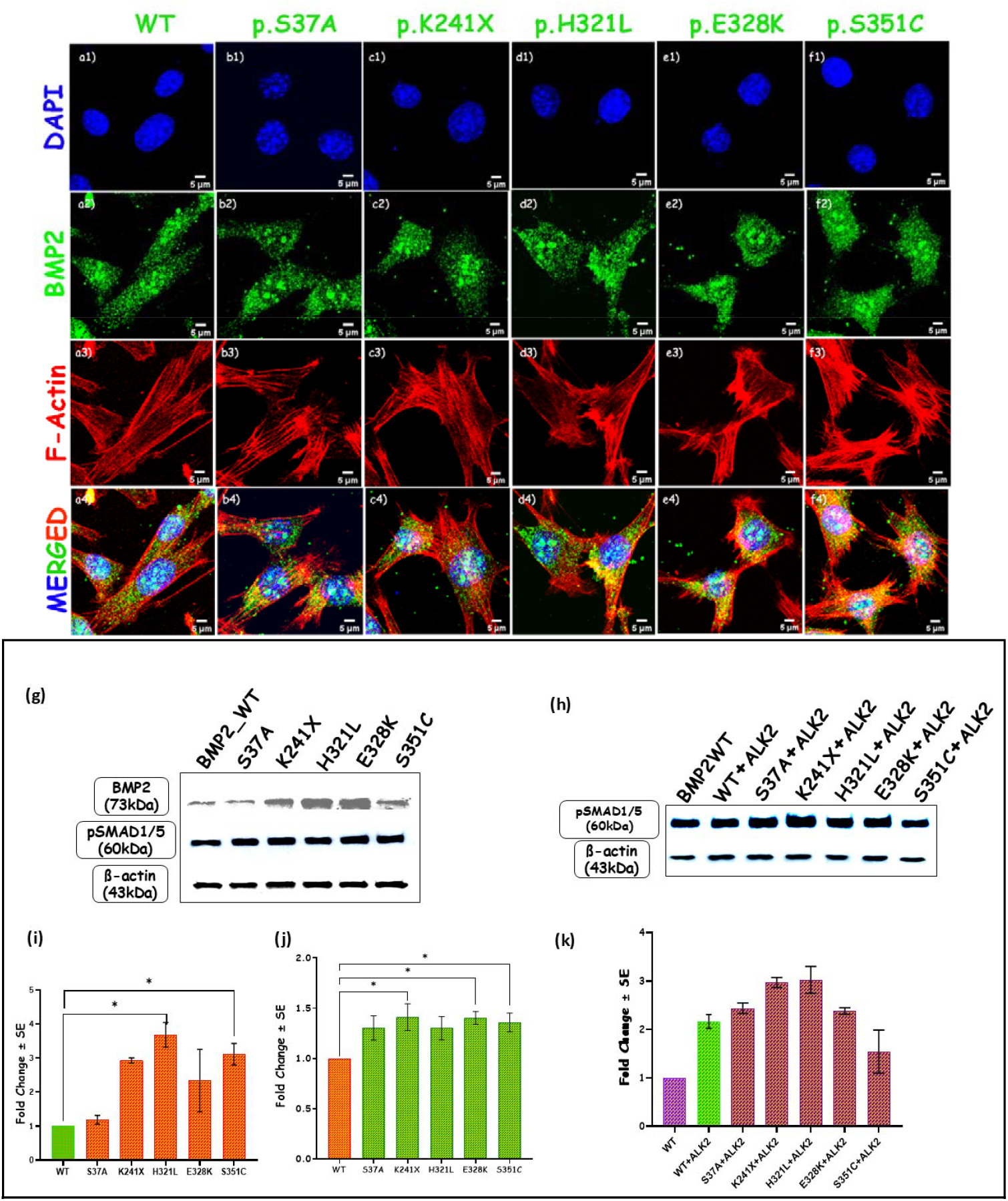
(i) Expression analysis by immunostaining. (a1-f4) showing nuclear and cytoplasmic localization of *BMP2 WT* and muteins (p.Ser37Ala, p.Lys241X, p.His321Leu, p.Glu328Lys and p.Ser351Cys) in H9c2 cells, (ii) expression analysis by western blotting of *BMP2 WT* and MUT in P19 cells (g) expression level of BMP2 and pSmad1/5 due to WT and all five variants with graphical representation in terms of fold change (i & j), (h) expression level of pSmad1/5 due to WT and all five variants (p.Ser37Ala, p.Lys241X, p.His321Leu, p.Glu328Lys and p.Ser351Cys) when co-transfected with *ALK2* with graphical representation in terms of fold change (k).

### 3.8 Effect of variants on the level of expression of BMP2 and pSMAD1/5 protein

The expression level of BMP2 proteins due to all the five variants were analyzed in P19 cells. There was a significant increase in the expression level of BMP2 proteins due to two mature domain p.His321Leu (3.678-fold; p=0.0159) and p.Ser351Cys (3.108-fold; p=0.0451) variants (Fig. 3(ii-g). However, notable increase in the expression of BMP2 in response to p.Ser37Ala (1.180-fold; p=0.9974), p.Lys241X (2.926-fold; p=0.0644), and p.Glu328Lys (2.335-fold; p=0.2139) was reported [Fig. 3(ii-g)]. Additionally, there was significant increase in the phosphorylation of SMAD1/5 by 1.411-fold (p=0.0207), 1.402-fold; (p=0.0228), 1.358-fold (p=0.0379) in response to p.Lys241X, p.Glu328Lys, and p.Ser351Cys variants respectively [Fig. 4(ii-g)]. On the other hand, remarkable increment was observed in the expression of BMP2 protein by 1.304-fold (p=0.0727), and 1.300-fold (p=0.0755) due to p.Ser37Ala, and p.His321Leu. In co-transfection experiment with *ALK2*, the phosphorylation of SMAD1/5 was also found to be boosted due to p.Ser37Ala, p.Lys241X, p.His321Leu, and p.Glu328Lys variants by 1.126 (p= 0.8674), 1.371 (p= 0.1218), 1.396 (p= 0.0966), and 1.099 (p= 0.9444) fold respectively [Fig. 4(ii-h)]. Conversely, a slight decrease in the phosphorylation of SMAD1/5 caused by variant p.Ser351Cys by 1.404 fold (p= 0.2684) was noted [Fig. 4(ii-h)].

### 3.9 Impact of variants on the transcriptional activity of downstream promoters

Dual reporter luciferase assay was performed to estimate the functional efficiency of BMP2 muteins on the transcriptional activity of well-known promoter of BMP signaling i.e., *Id1-luc, Id3-luc, p(SBE)*_*4*_*-luc*, and *Tlx2-luc*. The synergistic activity of *BMP2* with *ALK2* was also calculated by co-transfection experiment with each mentioned promoter. The transcriptional activity of all the promoters (*Id1-luc, Id3-luc, p(SBE)*_*4*_*-luc, and Tlx2-luc)* in response to either *BMP2* WT or mutants were normalized with the basal activity of these promoters. *BMP2* WT transactivated *Id1-luc* by 3.615-fold (p=0.009) (Fig. 5a), *Id3-luc* by 5.303-fold (p<0.0001) (Fig. 5c), *p(SBE)*_*4*_*-luc* by 3.033-fold (p=0.0075) (Fig. 5e) and *Tlx2-luc* by 2.905-fold (p=0.0156) (Fig. 5g). The impact of *BMP2* mutants on the transcriptional activity of these promoters was calculated as fold change by comparing with the WT. The pro-peptide domain variant p.Ser37Ala showed a significant reduction in transcriptional activity of promoter *Id1-luc* by 1.38-fold (p=0.3728) (Fig. 5a), *Id3-luc* by 2.18-fold (p=0.0015) (Fig. 5c), *p(SBE)*_*4*_*-luc* by 1.47-fold (p=0.3025) (Fig 4e), and *Tlx2-luc* by 1.60-fold (p=0.0245) (Fig 5g). Contrastingly, the stop codon variant p.Lys241X significantly enhanced the activity of promoters *Id1-luc* by 2.24-fold (p= 0.0004) (Fig. 5a), *Id3-luc* by 1.433-fold (p= 0.0089) (Fig. 5c), *p(SBE)*_*4*_*-luc* by 2.06-fold (p= 0.0001) (Fig. 5e) and *Tlx2-luc* 2.11-fold (p= 0.0007) (Fig. 5g). Similarly, the transcriptional activity of the promoters *Id1-luc, Id3-luc, p(SBE)*_*4*_*-luc*, and *Tlx2-luc* was significantly increased in response to other three mature peptide domain variants namely p.His321Leu, p.Glu328Lys and p.Ser351Cys. In case of p.His321Leu, the activities of *Id1-luc, Id3-luc, p(SBE)*_*4*_*-luc*, and *Tlx2-luc* promoters were increased by 1.92-fold (p=0.0023) (Fig. 5a), 1.89-fold (p <0.0001) (Fig. 5c), 1.74-fold (p=0.0036) (Fig. 5e) and 1.61-fold (p= 0.0222) (Fig. 5g) respectively. Correspondingly, significant enhanced transactivation of promoters *Id1-luc* by 2.12-fold (p <0.0007) (Fig. 5a), *Id3-luc* by 1.58-fold (p <0.0008) (Fig. 5c), *p(SBE)*_*4*_*-luc* by 2.33-fold (p <0.0001) (Fig. 5e), and *Tlx2-luc* by 3.21-fold (p= 0.0008) (Fig. 5g) was observed due to another mature peptide domain variants p.Glu328Lys. Another mature peptide domain variant p.Ser351Cys also significantly elevated the promoter activity of *Id1-luc* by 1.60-fold (p= 0.0226) (Fig. 5a), *Id3-luc* by 1.66-fold (p= 0.0003) (Fig. 5c), *p(SBE)*_*4*_*-luc* by 1.81-fold (p= 0.0016) (Fig. 5e), and *Tlx2-luc* by 1.60-fold (p= 0.0245) (Fig. 5g).

**Fig. 5.**
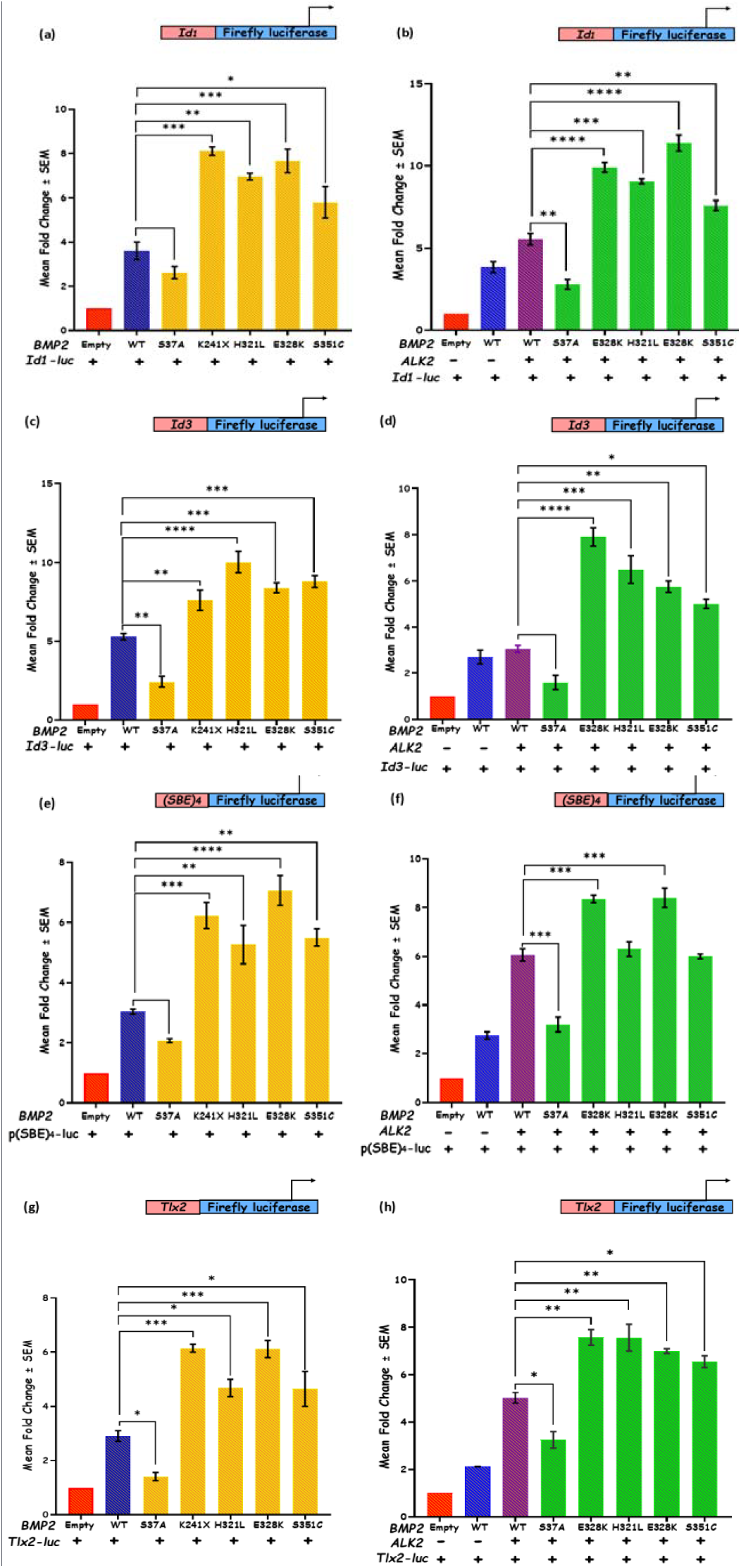
Transactivation assay by dual luciferase for measuring the transcriptional activity of *Id1-luc, Id3-luc, Tlx2-luc and (SBE)*_*4*_*-luc* Activation promoters. (a) of *Id1-luc* in response to *BMP2 WT* and MUT alone. (b) Synergistic activity of *Id1-luc* in response to WT and MUT when co-transfected with *ALK2*. (c) Luciferase result showing activity of *Id3-luc* in response to *BMP2 WT* and MUT alone. (d) Transactivation assay result indicating the synergistic effect *BMP2 WT* and MUT on transcriptional activity of *Id3-luc* when co-transfected with *ALK2*. (e) Transcriptional activity of *(SBE)*_*4*_*-luc* due to *BMP2 WT* and MUT alone. (f) Luciferase assay showing synergistic activity of *(SBE)*_*4*_*-luc* due to WT and MUT when co-transfected with *ALK2*. (g) Transcriptional activity of *Tlx2-luc* due to *BMP2 WT* and MUT alone (h) Synergistic effect of *BMP2 WT* and MUT on transcriptional activity of *Tlx2-luc* when co-transfected with *ALK2*.

Furthermore, when *BMP2* WT was co-transfected with *ALK2*, it synergistically trans-activated *Id1-luc* by 1.45-fold (p=0.0217) (Fig. 5b), *Id3-luc* by 1.13-fold (p=0. 0.9431) (Fig. 5d), *p(SBE)*_*4*_*-luc* by 2.2-fold (p<0.0001) (Fig. 5f), and *Tlx2-luc* 2.36-fold (p=0.0005) (Fig. 5h). The synergistic activity of the all four promoters was also decreased by p.Ser37Ala variant when co-transfected with *ALK2*, such as *Id1-luc* by 1.98-fold (p= 0.0012) (Fig. 5b), *Id3-luc* by 1.91-fold (p= 0.0575) (Fig. 5d), *p(SBE)*_*4*_*-luc* by 1.89-fold (p= 0.0002) (Fig. 5f), and *Tlx2-luc* by 1.55-fold (p= 0.0123) (Fig. 5h), which is comparable to the effect seen by p.Ser37Ala alone. In contrast, all four distal variants (p.Lys241X, p.His321Leu, p.Glu328Lys and p.Ser351Cys) potentiated the activity of all the promoters when co-transfected with *ALK2*. Interestingly, the variant p.Lys241X escalated the activity of *Id1-luc, Id3-luc, p(SBE)*_*4*_*-luc*, and *Tlx2-luc* by 1.78-fold (p <0.0001) (Fig. 5b), 2.59-fold (p <0.0001) (Fig. 5d), 1.38-fold (p= 0.0007) (Fig. 5f) and 1.51-fold (p= 0.0013) (Fig. 5h) respectively. Moreover, the synergistic activity of *ALK2* also significantly elevated the promoter activities of *Id1-luc* by 1.63-fold (p=0.0003) (Fig. 5b), *Id3-luc* by 2.13-fold (p=0.0004) (Fig. 5d), *p(SBE)*_*4*_*-luc* by 1.04-fold (p=0.9525) (Fig. 5f), and *Tlx2-luc* by 1.51-fold (p=0.0013) (Fig. 5h) by variant p.His321Leu. Likewise, the combinatorial effect with *ALK2* also increased the transcriptional activity of promoters *Id1-luc, Id3-luc, p(SBE)*_*4*_*-luc*, and *Tlx2-luc* by 2.05-fold (p<0.0001) (Fig. 5b), 1.89-fold (p= 0.0018) (Fig. 5d), 1.39-fold (p= 0.0006) (Fig. 5f) and 1.39-fold (p= 0.0066) (Fig. 5h) respectively in response to variant p.Glu328Lys. Similarly, the variant p.Ser351Cys, in combination with *ALK2* enhance the activities of promoter i.e *Id1-luc, Id3-luc, p(SBE)*_*4*_*-luc*, and *Tlx2-luc*, by 1.37-fold (p= 0.0079) (Fig. 5b), 1.63-fold (p= 0.0128) (Fig. 5d), 1.00-fold (p= 0.9998) (Fig. 5f) and 1.30-fold (p= 0.0278) respectively (Fig. 5h).

### 3.10 Effect of BMP2 variants on the expression of its downstream targets and other co-transcriptional activators

The qRT-PCR assay was carried out to analyse the effect of *BMP2* mutants on the expression of cardiac enrich transcription factors and co-transcriptional activators. All the five variants showed significant enhanced expression of downstream targets of BMP signaling. A significant increase in the expression of *Bmp2* in response to all the five p.Ser37Ala (1.393-fold, p<0.0001); p.Lys241X (1.448-fold, p<0.0001); p.His321Leu (1.299-fold, p=0.0023); p.Glu328Lys (1.544-fold, p<0.0001); p.Ser351Cys (1.402-fold, p<0.0001) variants were demonstrated (Fig. 6a). Further, the expression of *Bmp7* (form a potent heterodimer with *Bmp2*) was evaluated, a significant increment in the expression induced by four p.Ser37Ala (1.487-fold, p<0.0001); p.His321Leu (1.372-fold, p=0.0005); p.Glu328Lys (1.454-fold, p<0.0001); p.Ser351Cys (1.524-fold, p<0.0001) variants was noted while p.Lys241X (1.201-fold, p=0.1040) variant slightly increased the expression of *Bmp7* (Fig. 6b). Moreover, the expression of another transcription factor *Nkx2*.*5* was also significantly increased by p.Ser37Ala (1.347-fold, p<0.0001); p.Lys241X (1.187-fold, p<0.0134); p.His321Leu (1.25-fold, p=0.0004); p.Glu328Lys (1.243-fold, p=0.0006); p.Ser351Cys (1.492-fold, p<0.0001) (Fig. 6c). Besides, the up-regulation of *Gata4* was also significantly notable by 1.280-fold (p=0.0004), 1.432-fold (p<0.0001), 1.282-fold (p<0.0003), 1.280-fold (p=0.0004), 1.244-fold (p=0.0042) in response to variants p.Ser37Ala, p.Lys241X, p.His321Leu, p.Glu328Lys and p.Ser351Cys respectively (Fig. 6d). Furthermore, all the five mutants p.Ser37Ala (1.347-fold, p<0.0001), p.Lys241X (1.254-fold, p=0.0078), p.His321Leu (1.21-fold, p=0.0493), p.Glu328Lys (1.273-fold, p=0.0034), and p.Ser351Cys (1.359-fold, p<0.0001) caused significant enhanced expression of *Irx4* (Fig. 6e).

**Fig. 6.**
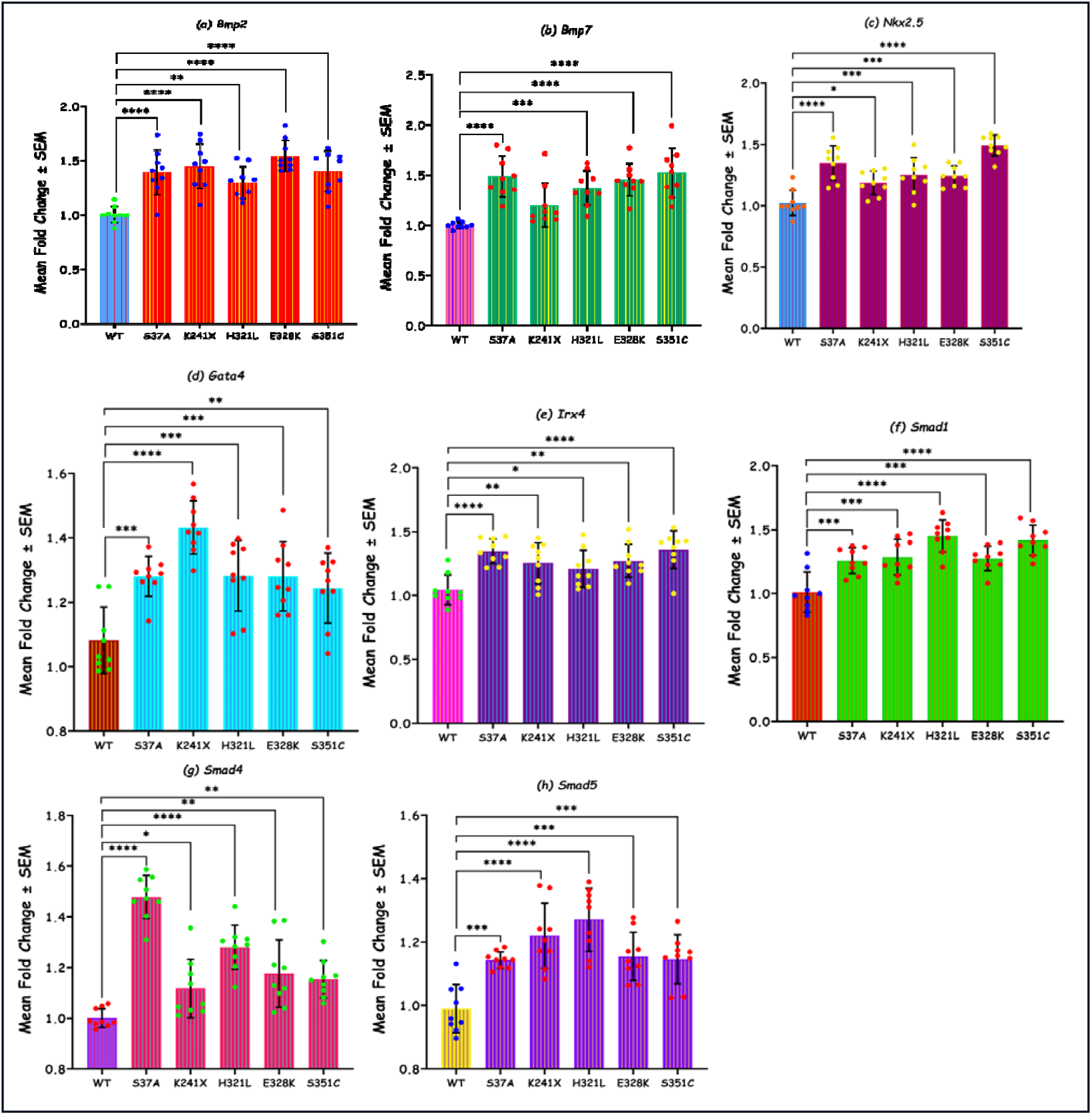
Quantitative assessment by real time PCR showing the expression of different downstream cardiac targets genes of *BMP2* (a) *Bmp2*, (b) *Bmp7*, (c) *Nkx2*.*5*, (d) *Gata4*, (e) *Irx4*, (f) *Smad1*, (g) *Smad4*, (h) *Smad5*, in response to all the five variants (p.Ser37Ala, p.Lys241X, p.His321Leu, p.Glu328Lys and p.Ser351Cys) of *BMP2*.

The increased expression caused by p.Ser37Ala (1.258-fold, p=0.0006), p.Lys241X (1.286-fold, p=0.0001), p.His321Leu (1.451-fold, p<0.0001), p.GlLys328K (1.275-fold, p=0.0002) and p.Ser351Cys (1.418-fold, p<0.0001) variants for co-transcriptional activator *Smad1* was also significantly acceptable (Fig. 6f). The effect on the up-regulation of *Smad4* was significant due to variants p.Ser37Ala (1.478-fold, p<0.0001), p.Lys241X (1.117-fold, p=0.0456), p.His321Leu (1.28-fold, p<0.0001), p.Glu328Lys (1.176-fold, p=0.0011) and p.Ser351Cys (1.154-fold, p=0.0049) (Fig. 6g). Additionally, the upregulation of another co-transcriptional activator *Smad5* due to these mutants p.Ser37Ala (1.143-fold, p=0.0008), p.Lys241X (1.22-fold, p<0.0001), p.His321Leu (1.27-fold, p<0.0001), p.Glu328Lys (1.155-fold, p=0.0003) and p.Ser351Cys (1.146-fold, p=0.0007) was also significant (Fig. 6h).

## 4. Discussion

*BMP2* is one of the well-studied members of TGF-β superfamily, known to play critical role in embryogenesis. Like other member of BMPs, BMP2 is a secretory signaling protein (42.5KDa), encoded by 396 amino acids (aa), secreted as precursor proteins with an N-terminal signal peptide (23 aa), a pro-domain (223 aa) for folding and secretion, and a C-terminal mature peptide domain (113 aa). The signal peptide facilitates secretion of the mature peptide via endoplasmic reticulum (von Heijne 1990). Precursors of BMP2 are formed in the cytoplasm as dimeric pro-protein complexes, which are cleaved by pro-protein convertases to generate N– and C-terminal fragments. The C-terminal mature fragments are capable of binding to its receptor, while the non-covalently associated, already cleaved pro-domains, play an important regulatory role by promoting the stability and dimerization of mature peptides (Moustakas and Heldin 2009; Sedlmeier and Sleeman 2017). The mature domain forms homo/heterodimers (BMP2, BMP7) by di-sulphide bond and secreted out into extracellular matrix for the commencement of BMP signaling (Hogan 1996). The secreted homo/heterodimers bind to the hetero-tetrameric receptor complexes which get activated and phosphorylate downstream, SMAD transcription modulators to transduce the BMP signal to the nucleus for inducing the downstream target genes (Massague 2000) and regulate many aspects of skeletal, cardiovascular, craniofacial, and limb development (Wu et al. 2007).

*BMP2* gene is located on chromosome 20p12.3 (GRCh38/hg38 assembly; HGNC:1069). It comprises four exons and three introns, however the protein is encoded by exon 2 and 3 only. Monoallelic mutation in *BMP2* gene, exhibited multisystem malformations characterized by short stature, skeletal and craniofacial anomalies, eye and auditory defect as well as cardiac structural and rhythm defects (Lalani et al. 2009; Sahoo et al. 2011; Williams et al. 2012; Tan et al. 2017; Su et al. 2021; Priestley et al. 2023). However, many of these studies deal with either sub-microscopic deletions, duplications, frameshift or truncation mutations on the chromosomal segment (20p12.3) harbouring BMP2 gene. BMP2 deficient mice are embryonic lethal showing cardiac defects. Multiple studies have shown that expression of BMP2 is essential for endocardial cushion (EC) formation. Conditional deletion of *BMP2* in atrioventricular (AV) myocardium confirm the crucial role of BMP2 in EC EMT, as well as in formation of AV cushion (Ma et al. 2005). Considering the pivotal role of BMP2 on cardiac development, herein we investigate whether impaired BMP2 function, could cause CHD. In the present study, we have identified 5 nonsynonymous variations namely, p.Ser37Ala (in between signal peptide and pro-domain), p.Lys241X (in the pro-domain) and p.His321Leu, p.Glu328Lys and; p.Ser351Cys (in the mature domain) in 8 isolated cases of CHD, depicting an overall mutation frequency of 2.8% in our cohort (Indian population). The variant p.Ser37Ala (rs2273073) is already reported, which is linked to complex CHD phenotype (dTGA, and ASD, VSD, PA) in two unrelated probands. A non-sense variant (p.Lys241X) is confined to pro-peptide domain, identified in a patient with PTA with VSD phenotype. The mature domain harbour three other variants viz., p.His321Leu, p.Glu328Lys and p.Ser351Cys which are associated with dTGA, VSD and ToF & ASD (Pentalogy of Fallot; PoF) respectively (Table 1). The low allele frequencies (MAF>0.01) points to the pathogenic potential of these variants. In the present study, most of the phenotypes linked to *BMP2* variants are outflow tract and septal defects, which are presumed due to dysfunction of BMP2 during cardiac jelly formation and EMT. In our study cohort, we have recruited isolated CHD cases only (without any extracardiac phenotype), using strict inclusion and exclusion criteria, thereby the influence of defective BMP2 signaling on the formation/ malformation of heart could be explicitly assessed. To date, only a limited number of studies have elucidated the association of single nucleotide rare variations (SNVs) in *BMP2*, with non-syndromic CHD and functional analysis of such variants remain inadequate. Earlier in a study by Li et al, (2016) has provided evidence for the association of two reported SNPs (rs1049007, c.584 A>G and rs235768, c.893 A>T) in *BMP2* with CHD with statistical significance in Chinese population (Li et al. 2016). Later, another case-control study from Greece, using 52 CHD cases and 100 control blood samples has also reported the same SNPs, however without any significant association (Bobos et al. 2023a). Further no significant differential expression of BMP2 and BMP4 could be detected in the cardiac tissue biopsies from the same patients-control group. Nonetheless, Ahluwalia and Gelb (2021) identified a nonsense variant (p.R105X) in *BMP2*, in a 3 years old male child, detected with bicuspid aortic valve (BAV), however, functional characterization of the variant has not been carried out (Ahluwalia and Gelb 2021). Another study by Yogi and his colleagues have reported a novel pathogenic frameshift variant c.231dup (p.Tyr78Leufs*38) in *BMP2* which is associated with isolated dextrocardia, *situs solitus* in the second twin of a familial case (Yogi et al. 2023). In human, the expression of BMP2 is abundant with quantitatively similar expression in CHD-affected and normal hearts. However, sustained expression of mRNA and protein is observed in CHD individuals (Bobos et al. 2023b). In addition to CHD, other multisystem pathogenic conditions are also reported often with syndromic manifestation, in patients harbouring *BMP2* mutations. A case-control study reveal two pathogenic SNPs (p.Ser37Ala and p.Ala190Ser) identified in *BMP2* contribute to the susceptibility to congenital combined pituitary hormone insufficiency (CPHD) (Breitfeld et al. 2013). We have also identified the p.Ser37Ala variant in two unrelated isolated CHD cases. However, pituitary deficiency is not recognized in any of the CHD cases, in our study. A collaborative study by Tan et al, (2017) showed that monoallelic truncating and frameshift *BMP2* variants and deletions are associated with short stature, a recognizable craniofacial gestalt, skeletal anomalies, and congenital heart disease in 12 individuals from eight unrelated families (Tan et al. 2017). In a recent study, 7/18 patients showing missense variation, also exhibited skeletal and other extracardiac anomalies in addition to heart defects, which is similar to other patients carrying frameshift and nonsense mutations (Priestley et al. 2023). Many pathogenic variants have also been reported in other members of BMPs viz., *BMP4* (Posch et al. 2007; Qian et al. 2014; Li et al. 2016; Wang et al. 2024) and *BMP10* (Dong et al. 2024) which are shown to be associated with sporadic cases of non-syndromic CHD cases. Dispensable role of BMP4 for skeleto-genesis and bone repair has been demonstrated in mouse model (Wang et al. 2024). Conversely, recombinant BMP2 is shown to induce cartilage and bone ectopically (Wang et al. 2024). Different downstream members of BMP signaling also contribute to the pathogenesis of CHD namely *ALK2* (Joziasse et al. 2011), *ALK3* (Demal et al. 2019), *BMPRII* (Roberts et al. 2004; Qi et al. 2020), *SMAD1* (Wang et al. 2022), *SMAD4* (Wang et al. 2023), *GATA4* (Butler et al. 2010; Dixit et al. 2019), *NKX2*.*5* (McElhinney et al. 2003), *MEF2C* (Qiao et al. 2017; Lu et al. 2018), *IRX4* (Cheng et al. 2011). The null mice of *Id1*, a downstream effector of *Bmp2* exhibit valvular defects and outflow tract atresia as *Id1* is a crucial gene for endocardial EMT in both chick and mouse embryos (Katagiri et al. 2002). ID proteins are major players in early development and have crucial role in the differentiation and proliferation of cardiac progenitor cells and mature cardiomyocytes at multiple stages during heart development (Hu et al. 2019). Additionally, the double or triple Id knockout embryos (*Id1*/*Id2, Id2*/*Id3, Id1/Id3* or *Id1*/*Id2*/*Id3*) demonstrate severe cardiac defects including valvular and septal defects, outflow tract atresia, impaired ventricular trabeculation, thinning of the compact myocardium layers, and the embryos die at mid-gestation (Fraidenraich et al. 2004). In BMPR1a-cKO hearts, the expression of ID1/3 was absent from the embryonic atrioventricular canal (AVC) which affects the formation of the atrioventricular valves and adjacent septa (Kaneko et al. 2008). Further, mice model of Noggin^-/-^ (a dedicated BMP antagonist) and Smad6^-/-^ (Bmp-specific nuclear inhibitor), with increased of BMP function, exhibited thickened or hyperplastic valve (Conway et al. 2011).

Functional validation of all the *BMP2* variants by *in vitro* and *in silico* analyses, four out of five identified variants i.e., p.Lys241X in the pro-domain and three other mature domain variants p.His321Leu, p.Glu328Lys and p.Ser351Cys demonstrated gain-of-function (GoF), while the N-terminal variant, p.Ser37Ala, adjacent to signal peptide exhibited loss-of-function (LoF) activity. The phylogenetic analysis using multiple sequence alignment, illustrate that the amino acid residues (Lys241, His321, Glu328 and Ser351) those suffer mutation, are highly conserved, which suggest their essential role in protein structure and function. Further, ‘Varcard’ analysis of these 4 variants are predicted to be disease-causing and damaging. However, the p.Ser37Ala (already known) variant is speculated to be benign and tolerated. Our Western blot analysis revealed increase in the expression of mutant BMP2 proteins, by p.Lys241X, p.His321Leu, p.Glu328Lys and p.Ser351Cys variants except for variant p.Ser37Ala. However, when phosphorylation of downstream SMAD1/5 was studied overall the enhanced phosphorylation due to all the variants was observed. Smad phosphorylation is crucial for downstream signaling, that trigger crucial target gene expression inducing cellular differentiation and organ development (Dexheimer et al. 2016; Salazar et al. 2016; Lowery and Rosen 2018). Contrarily, our luciferase assay using four downstream gene’s promoters such as *Id1-luc, Id3-luc, Tlx2-luc, and p(SBE)*_*4*_*-luc* have revealed a decrease in the activity of all the promoters, by p.Ser37Ala variant, among these *Id3-luc* and *Tlx2-luc* showed significant depletion. Contrarily, significant upregulations in the transactivation of *Id1-luc, Id3-luc, Tlx2-luc, and p(SBE)*_*4*_*-luc* are observed in response to the non-sense (p.Lys241X) variant and the three mature domain (p.His321Leu, p.Glu328Lys and p.Ser351Cys) variants. It is very well established that BMP2 binds with BMPRI/ACVRI (ALK2) and BMPRII receptors, which subsequently trans-phosphorylate and activate SMAD1/5/8, thus inducing intracellular SMAD signaling, which further transduce the signal to nucleus to activate target gene promoters (Paul et al. 2005; Ye et al. 2023). Any perturbation in the cooperative binding of BMP2 with ALK2 would likely affect the spatial and temporal expression of the tightly regulated downstream signaling which is crucial for cardiogenesis. Therefore, we investigated the effect of *BMP2* variants on the synergistic transactivation of *Id1-luc, Id3-luc, Tlx2-luc, and p(SBE)*_*4*_*-luc* with *ALK2*. The combinatorial effect of *ALK2* also depicted a similar trend, i.e., a significant decline in the activity of all four promoters by p.Ser37Ala variant. The down regulation of all four promoters i.e., *Id1, Id3, Tlx2, and (SBE)*_*4*_, in response to Ser37Ala variant suggests impaired ER-mediated transport and secretion of the mutant protein. However, all other variants (p.Lys241X, p.His321Leu, p.Glu328Lys and p.Ser351Cys) displayed significantly enhanced transactivation of all the promoters irrespective of synergistic effect of *ALK2*, implicating the GoF activity. In a recent study, Ahluwalia and Gelb (2021) have identified a nonsense variant (p.R105X) in pro-peptide domain of *BMP2* in BAV which they assumed to be pathogenic due to haploinsufficiency of *BMP2*, albeit they have not performed any *in vitro* functional experiment (Ahluwalia and Gelb 2021). It has been described that mRNAs harbouring premature termination (nonsense) codons undergo a quality check and destroyed by nonsense-mediated mRNA decay (NMD), however if translated, these mRNAs can produce truncated proteins with dominant-negative or deleterious GoF activities (Chang et al. 2007). BMP2 pro-domain is reported to play a critical role in facilitating the cleavage and dimerization of mature domain not only of BMP2 homodimer itself but also dimerization of BMP6/7 (Chauhan et al., 2024). Further, BMP2 pro-domain interfere with binding of mature domain with its receptors thus inhibiting the downstream signaling (Hauburger, 2009). Therefore, it is speculated that a truncated protein produced here, by the nonsense variant become incompetent, thus enhancing receptor activity and downstream signaling consequently exhibiting deleterious GoF activity. Of note, the mature domain variants, p.His321Leu, p.Glu328Lys and p.Ser351Cys are localized in the convex hydrophobic segment of receptor binding (wrist and knuckle) region of mature protein (Zhang et al., 2021). Substitution of hydrophobic residues Leu, Lys probably enhance the binding efficiency with BMP receptors, thereby escalating the SMAD-mediated downstream signaling.

BMP2 is secreted either in the form of homodimer or heterodimer along with BMP7, prior to binding to cell surface receptors to initiate the canonical SMAD signaling. BMP2/BMP7 heterodimer is recognized to be biologically more active than either of homodimers (Suzuki et al. 1997; Aluganti Narasimhulu and Singla 2020). The overall increase in SMAD phosphorylation in all 5 variants, including p. Ser37Ala, albeit, the later showed reduced BMP2 protein level as well as decreased downstream promoter activity is intriguing. Nonetheless, when the expression pattern of *Bmp2* and *Bmp7* is checked by Real-time qRT-PCR, overexpression of both the *Bmp2 & Bmp7*, is observed with all five variations. Functional redundancy of BMP2, BMP4 and BMP7 which form heterodimer possibly contribute to increased SMAD signaling (Bandyopadhyay et al. 2006). Furthermore, qRT-PCR data also demonstrated enhanced expression of *Smad1, Smad4*, and *Smad5* in response to reported p.Ser37Ala as well as pro– and mature-domain variant (p.Lys241X, p.His321Leu, p.Glu328Lys and p.Ser351Cys). Interestingly, real time expression of *Nkx2*.*5, Gata4* and *Irx4* is also increased by all five variants. Bmp2-Smad1/5/8 signaling is known to play an important role during cardiogenesis by directly regulating transcription factors *Nkx2*.*5*, and *Gata4* (Schultheiss et al. 1997; Ladd et al. 1998; Prall et al. 2007; Si et al. 2014). In brief, above observations indicates the upregulation of BMP signaling, *Smad1* and *Smad5* are directly regulated by BMP-receptor complex which in turn form complex with *Smad4* to get translocated into nucleus to elicit the activity of multiple transcription factors (Morrell et al. 2016; Hu et al. 2019), which consequently control the expression of crucial regulators of cardiogenesis including *Nkx2*.*5, Gata4* and *Irx4* in P19 cells (Hu et al. 2019). Growing body of evidences also suggest that GoF mutation can lead to variety of developmental abnormality or diseases. Several studies have reported GoF mutations in BMP signalling causing abnormal gene function leading to disease phenotype. A Gof3 mutation in *ACVR1/ALK2* (a BMP receptor) is reported to cause fibrodysplasia ossificans progressive (Sanchez Duffhues et al.). Homozygous missense mutation in *BMPRIA*, causing elevated SMAD level, develop brachycephaly (Russell et al. 2019). Similarly, *ACVR1* having Gly328Trp or Gly328Glu mutation enhance SMAD1/5 phosphorylation, resulting in fibrodysplasia (Mucha et al. 2018). Interestingly, GoF mutation in *TGF-*β*1* (Yadav et al. 2022) and *CITED2* (Yadav et al. 2021) are also susceptible to CHD. Hence, the GoF activity *BMP2* variants, seen in the present study is likely responsible for the pathogenesis.

Although BMP2 like other members of BMP family is a secretory signaling molecule undergo ER-mediated cytoplasmic transportation to extracellular space, where the mature domain of BMP2 transduce SMAD-mediated downstream signaling. However, accumulation of BMP2 in the nucleus, as shown by our immunostaining is intriguing. A meticulously designed study by Felin et al., (2010) has demonstrated nuclear localization signal (NLS) sequence in BMP2 overlapping the site of proprotein processing, which generate alternative nuclear variant of BMP2 (nBMP2) that translocate to the nucleus instead of following secretory pathway. Though no direct DNA binding could be detected, indirect interaction via other protein is predicted. Hence, BMP2 signaling likely undergo leaky pathways where both secretory and nuclear transport co-exist. In this study, both WT and mutants show no difference in the level of nuclear localization. However, detailed investigation is required to understand the molecular mechanism involved.

Furthermore, a remarkable structural changes in mRNA of p.Lys241X, p.His321Leu, and p.Glu328Lys is noted by analysing base-pairing probabilities and accessibility profiles which led us to presume disrupt RNA folding, that may have impact on RNA stability, function and RNA-protein interactions which are crucial for the developmental genes like BMPs. This analysis underscores the impact of variants on RNA structure and function which provide insights into the molecular mechanisms related to disease pathogenesis (Salari et al. 2013). Likewise, secondary tertiary structure analysis of these variants disclosed the marked structural changes in muteins. A distinguished alterations in the helices and strands illustrated by variants p.His321Leu, p.Glu328Lys and p.Ser351Cys, which potentially have impact on protein folding and function (Supplementary Fig. 2). However, the termination of protein due to the non-sense variant (p.Lys241X) at pro-peptide region, and the substitution of histidine (positively charged) by leucine (neutral) for variant p.His321Leu, and glutamic acid (negatively charged) to lysine (positively charged) for variant p.Glu328Lys are speculated to change the overall net charge of BMP2 (Swencki-Underwood et al. 2008) that induces more secretion of mature BMP2 proteins and thereby upregulating the *SMAD* dependent pathway of BMP2. Altogether, integrating our *in silico* and *in vitro* experimental analysis, that demonstrate the upregulation of Smad signaling caused by GoF mutations in *BMP2*, which further reflect enhanced the secretion of BMP2 that consequently elevate a possible flawed BMP signaling. In addition, BMP signaling has been displayed to have cross-talk with other signaling networks during cardiogenesis like *Nodal, TGF-*β (Conway et al. 2011; Garside et al. 2013), *NOTCH* (Miyazono et al. 2005; Garside et al. 2013), *Wnt/* β –catenin etc. (Joziasse et al. 2008; Song et al. 2020) and increased BMP signaling possibly inhibiting other cardiac networks could results in such kind developmental abnormalities.

## 5. Conclusion

In short, here we identify five variations including a truncating nonsense variation in the pro-domain and three missense variations in the mature domain, in a large cohort of isolated CHD cases. Although majority of previous studies reported skeletal abnormalities, eye or auditory defects in addition to cardiac malformations, in patients carrying *BMP2* mutations, in the present study, our isolated CHD cohort exhibited exclusively cardiac septal and out flow track defects, corresponding to AV cushion glitch, predicted independent tissue-specific operation of BMP signaling network. Variable phenotypes observed in different study possibly due to tissue specific favourable/unfavourable interaction of different signaling network and tissue microenvironment culminating into such disease phenotypes. Furthermore, out of five variations detected in *BMP2*, four demonstrated GoF activity by upregulating downstream targets. Interestingly, in previous reports it has been demonstrated that cKO of *Bmp2, Bmp4* and *Bmpr2* hypomorph exhibited hypoplastic AV canal and OFT defects, while similar cKO of *Bmp4* and *Bmpr2* in second heart field and endothelium respectively caused hyperplastic thick valve, thus pleading the tissue specific influence of BMP singnaling and parallel influence of both GoF and LoF of BMP signaling (Conway et al., 2011). Additionally, in the present investigation, for the first time both *in vitro* and *in silico* functional analysis was performed, revealing strong upregulated BMP2 signaling, which consequently resulted in CHD pathogenesis that might pave the way in future for developing therapeutic strategies.

## List of abbreviations

CHD: Congenital Heart Disease
ASD: Atrial Septal defects
VSD: Ventricular Septal Defects
PDA: Patent Ductus Arteriosus
TGA: Transposition of Great Arteries
ToF: Tetralogy of Fallot
PoF: Pentalogy of Fallot
DORV: Double Outlet Right Ventricle
PFO: Patent Foramen Ovale
EMT: Epithelial to Mesenchymal Transformation

## Acknowledgment

We are deeply grateful to all the patients, their families, and the control individuals for their valuable participation in this study. We sincerely thank to Prof. T.K. Lahiri and Dr. Damyanti Agrawal, Department of Cardiovascular and Thoracic Surgery, IMS, BHU, Varanasi, for their constant support and encouragement during patient enrolment. We also acknowledge Dr. Ramkumar Sambasivan (InStem, Bengaluru, Karnataka, India) for providing P19 cells, Dr. Maria Genander (Karolinsk7a Institutet, Stockholm, Sweden) for the *Id1* luciferase reporter construct, and Dr. Daniel J. Bernard (Quebec, Canada) for the *Id3* luciferase reporter construct. We gratefully acknowledge the University Grants Commission (UGC), Government of India, for awarding Junior and Senior Research Fellowships (577/(CSIR-UGC NET JUNE 2018)) to Jyoti Maddhesiya.

## Data Availability

Raw data and derived data supporting the findings of this study are available from the corresponding author on request.

## CRediT authorship contribution

**Bhagyalaxmi Mohapatra:** Conceptualization, Data curation, Formal analysis, Funding acquisition, Investigation, Project administration, Resources, Supervision, Visualization, Writing – review & editing.

**Jyoti Maddhesiya**: Conceptualization, Data curation, Formal analysis, Methodology, Software, Validation, Visualization, Writing – original draft, Writing – review & editing.

**Ritu Dixit:** Methodology.

**Dharmendra Jain:** Investigation

