## Supplemental Information for "Implication of novel variants of *BMP2* in isolated congenital heart disease: Functional characterization by *in silico* and *invitro* approaches"

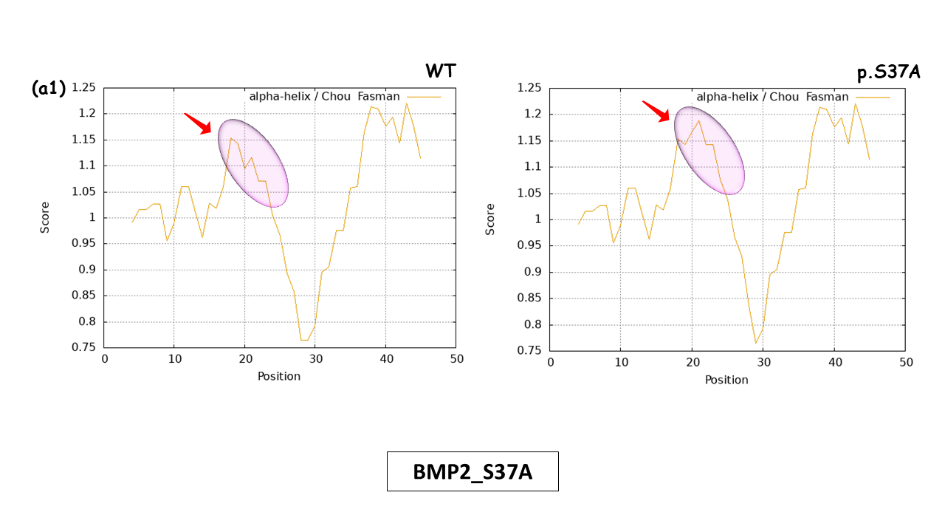

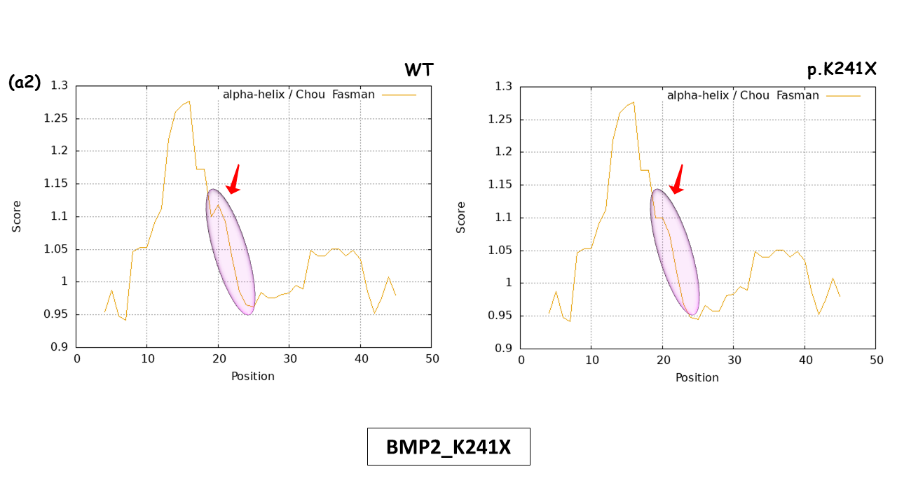

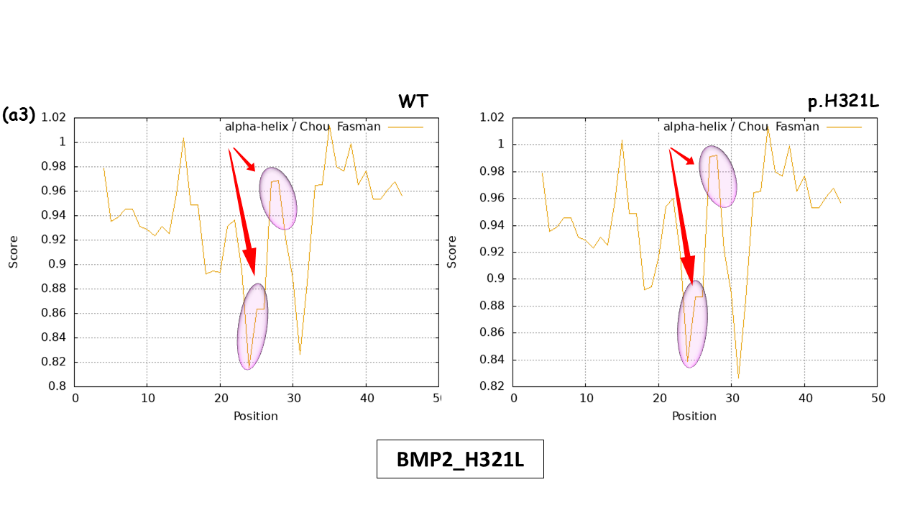

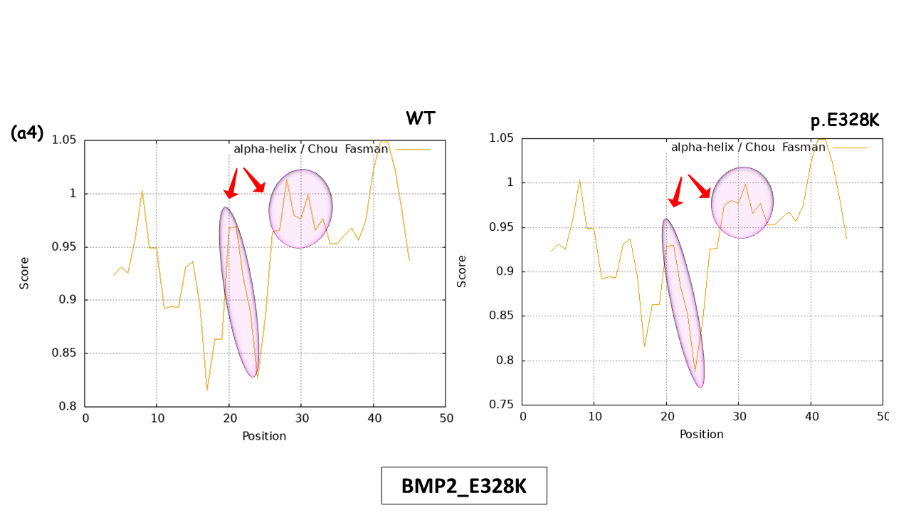

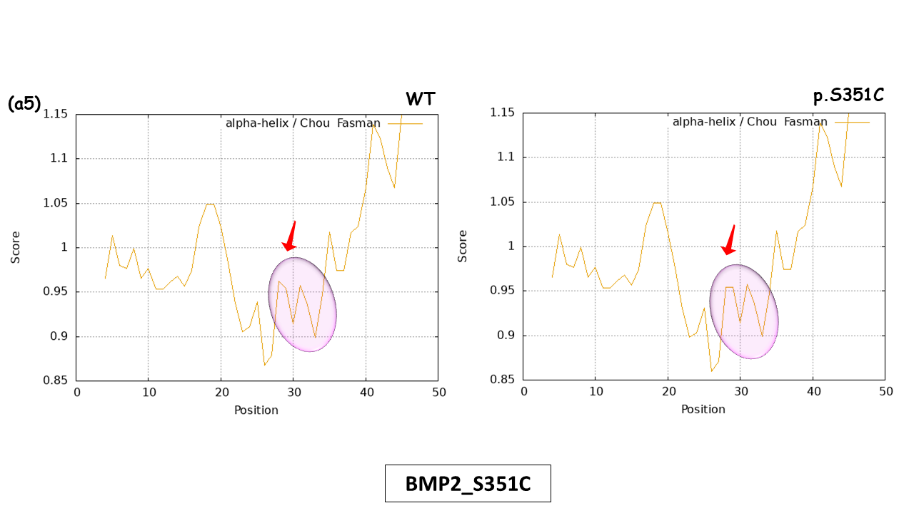

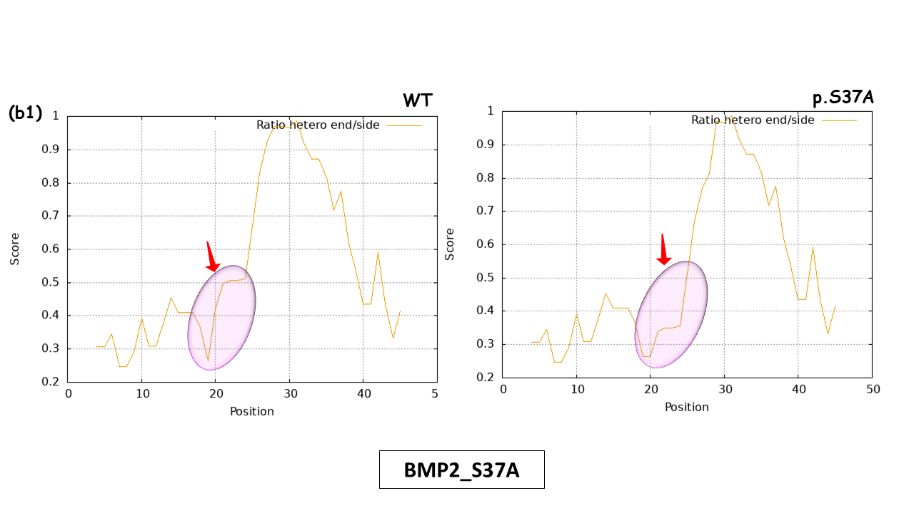

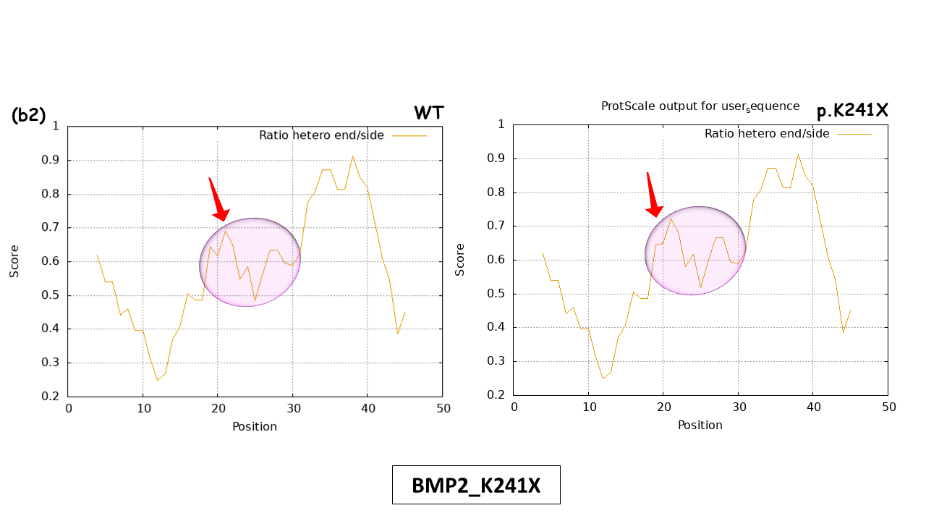

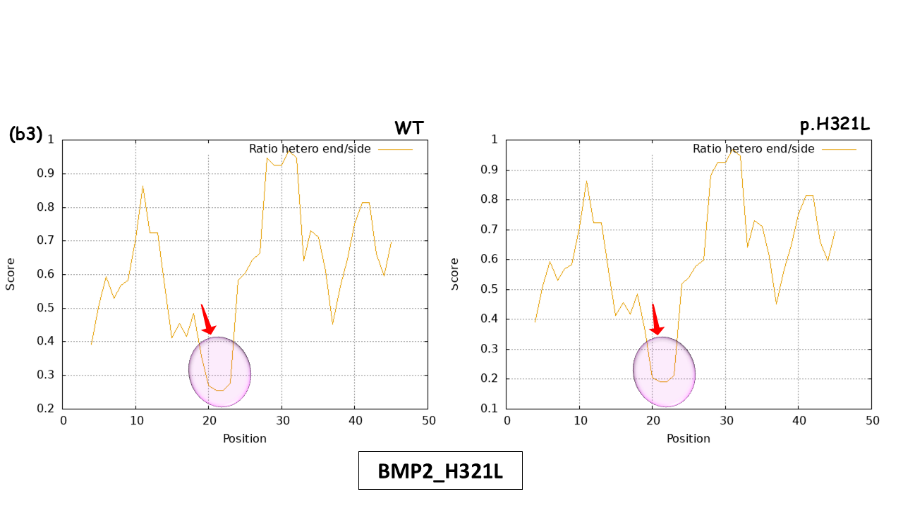

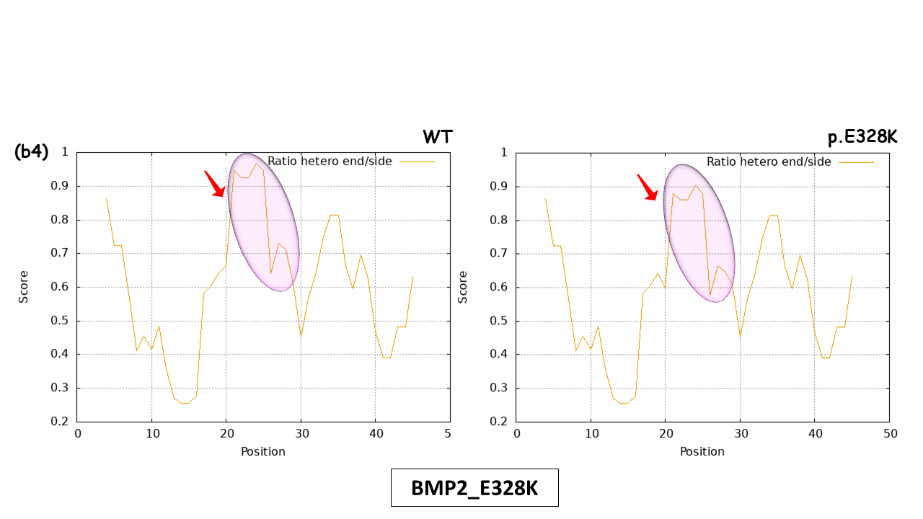

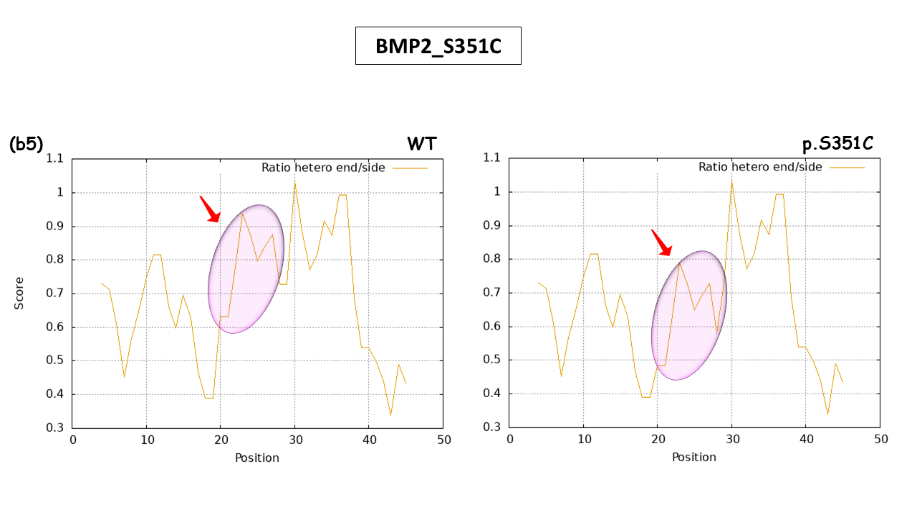

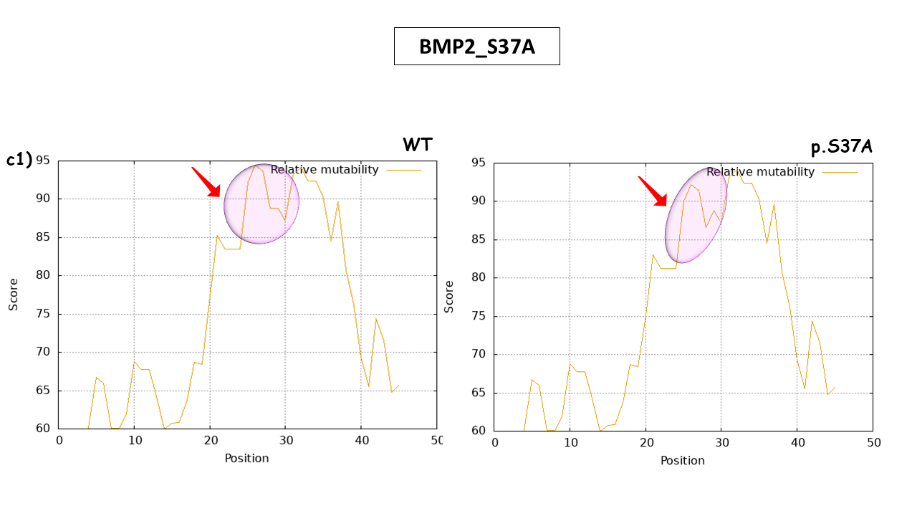

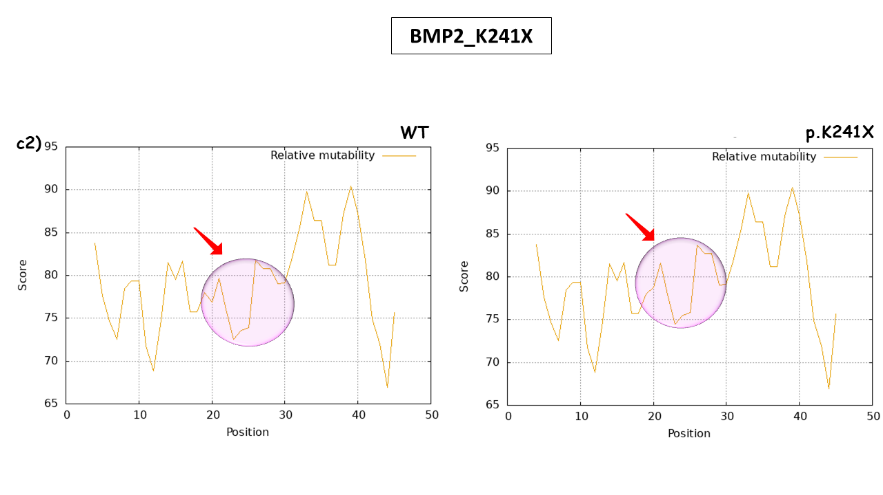

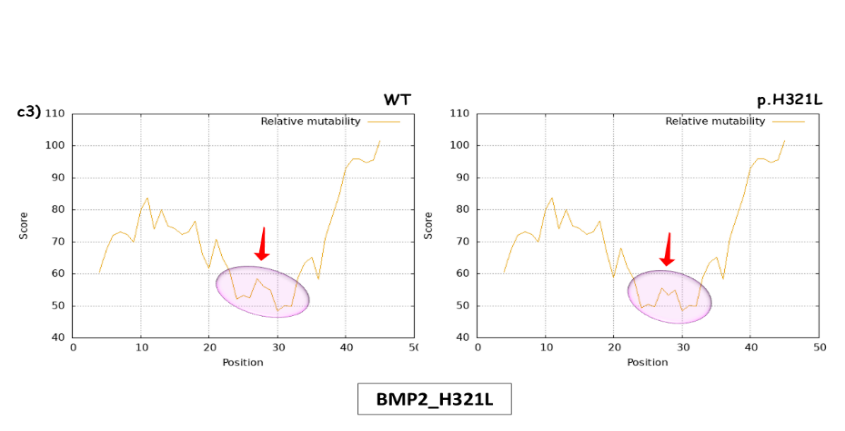

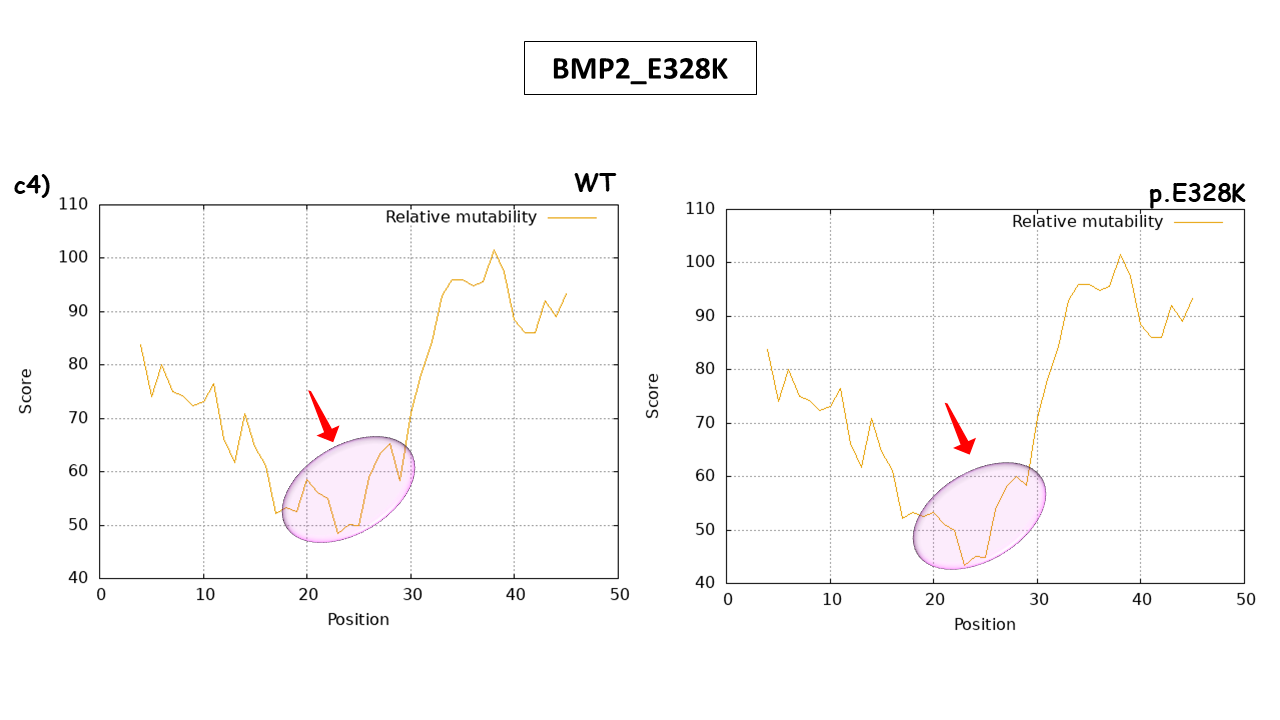

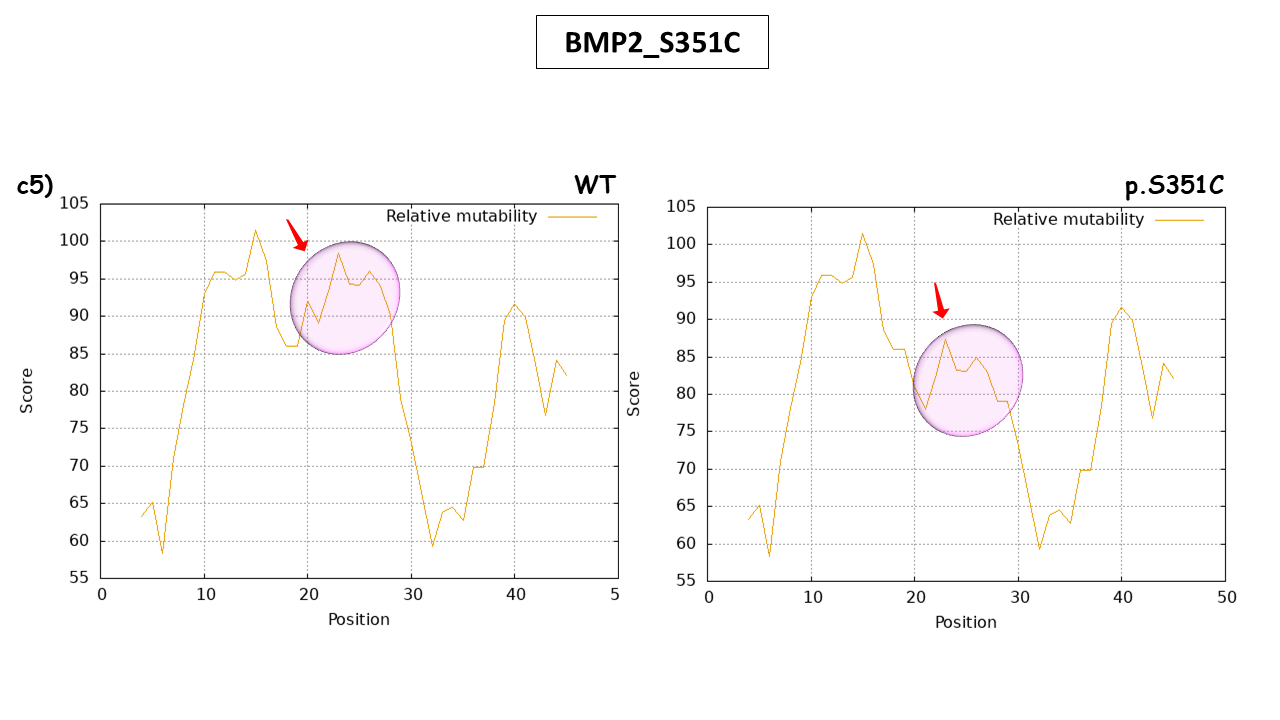

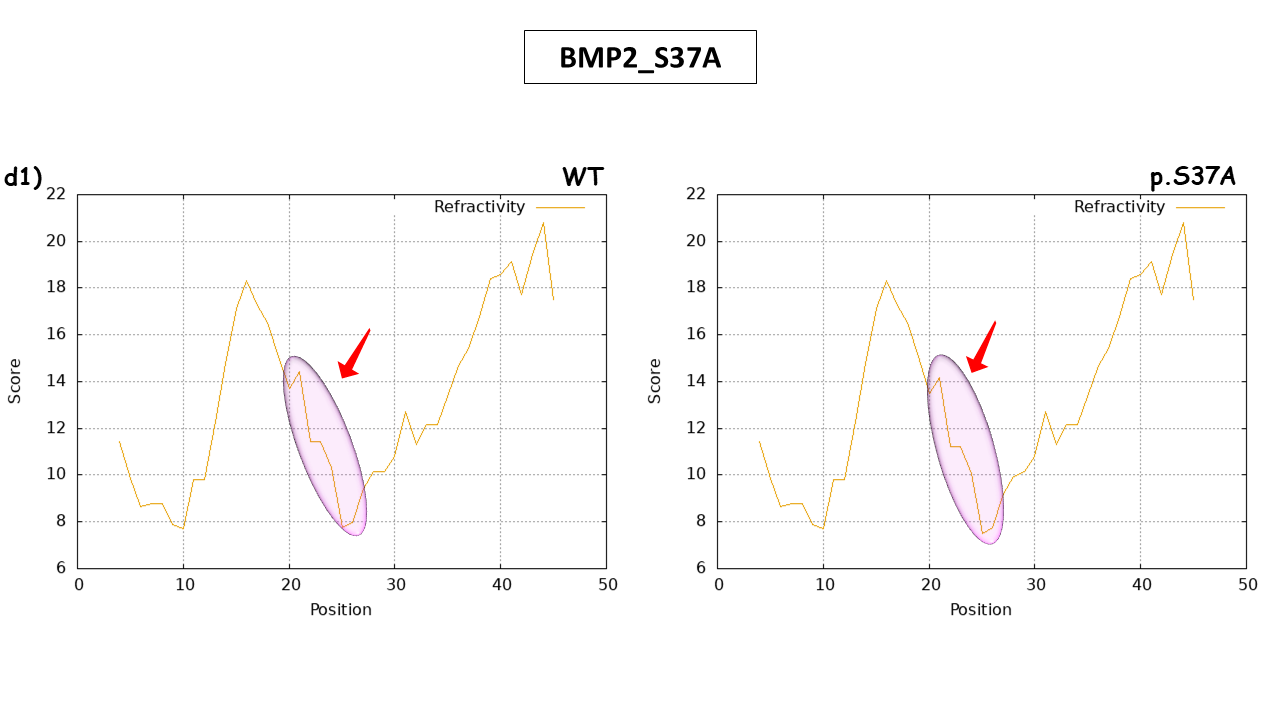

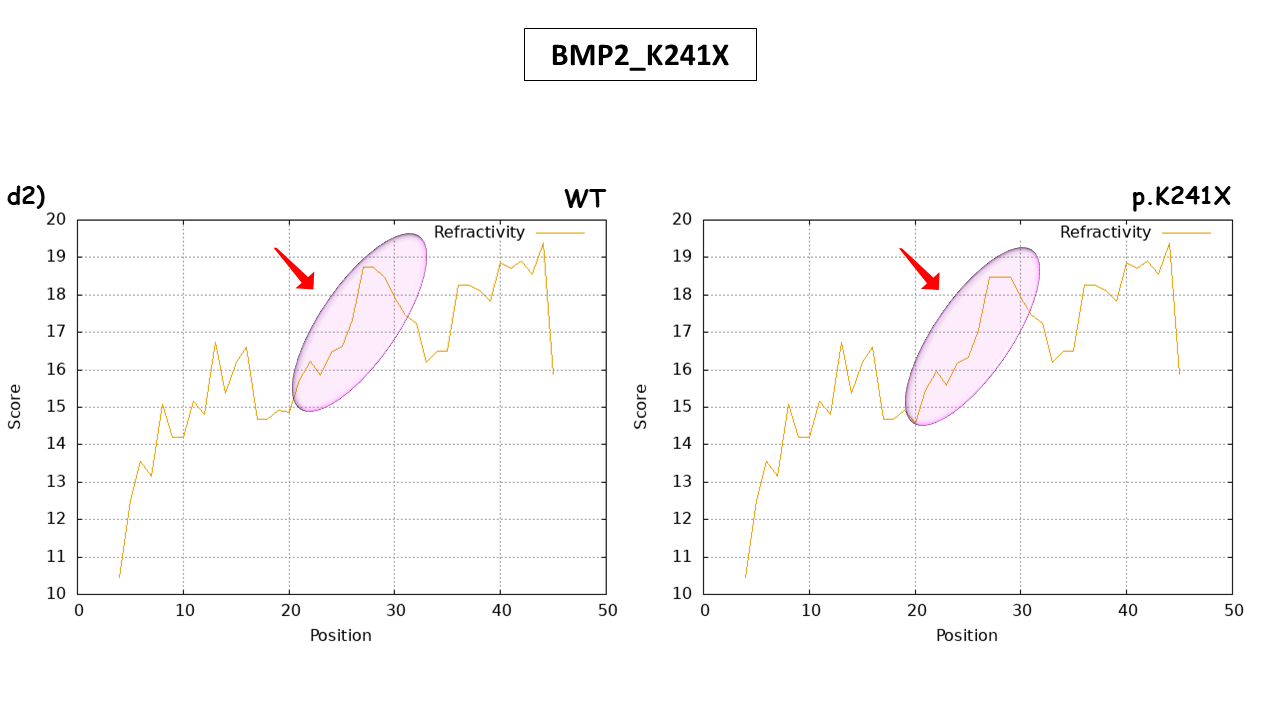

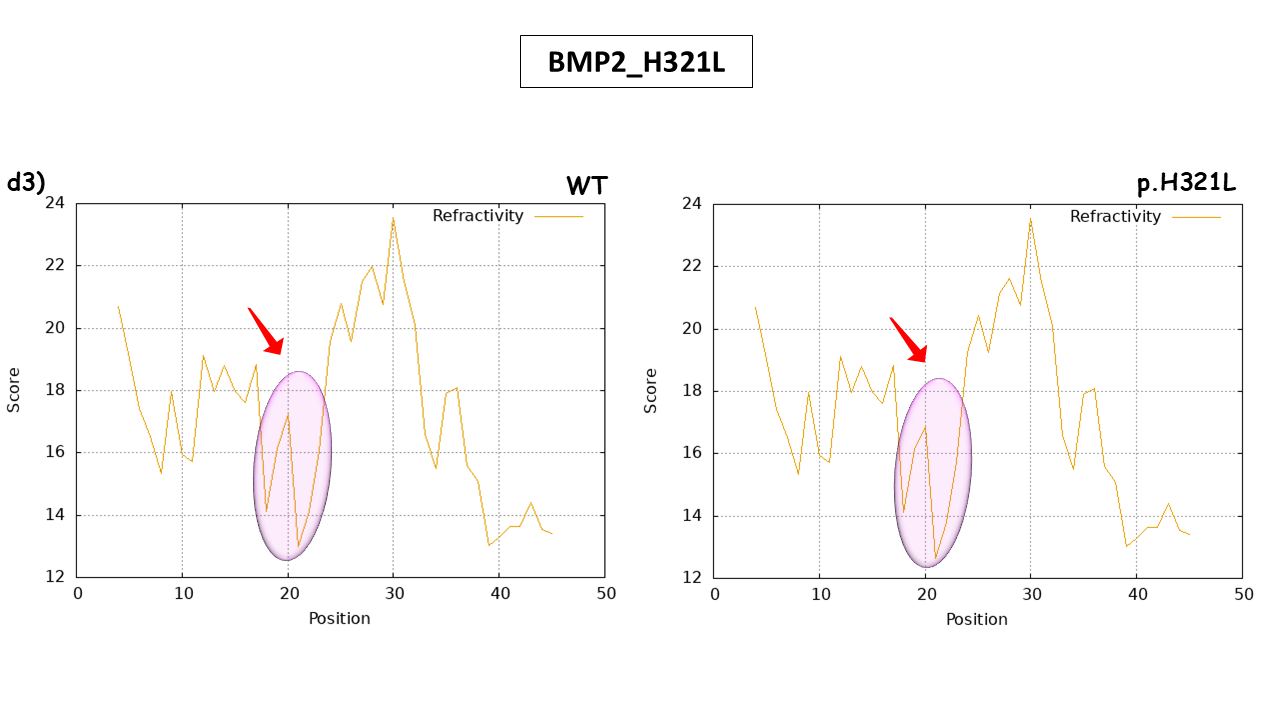

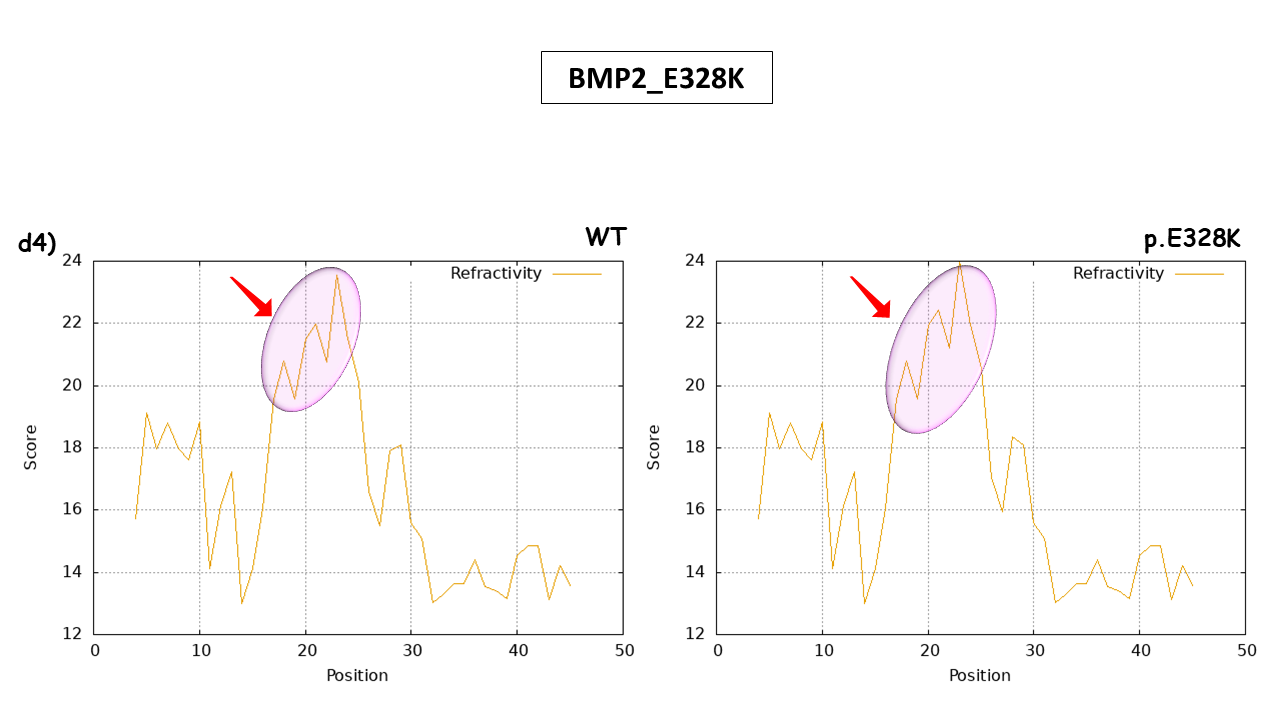

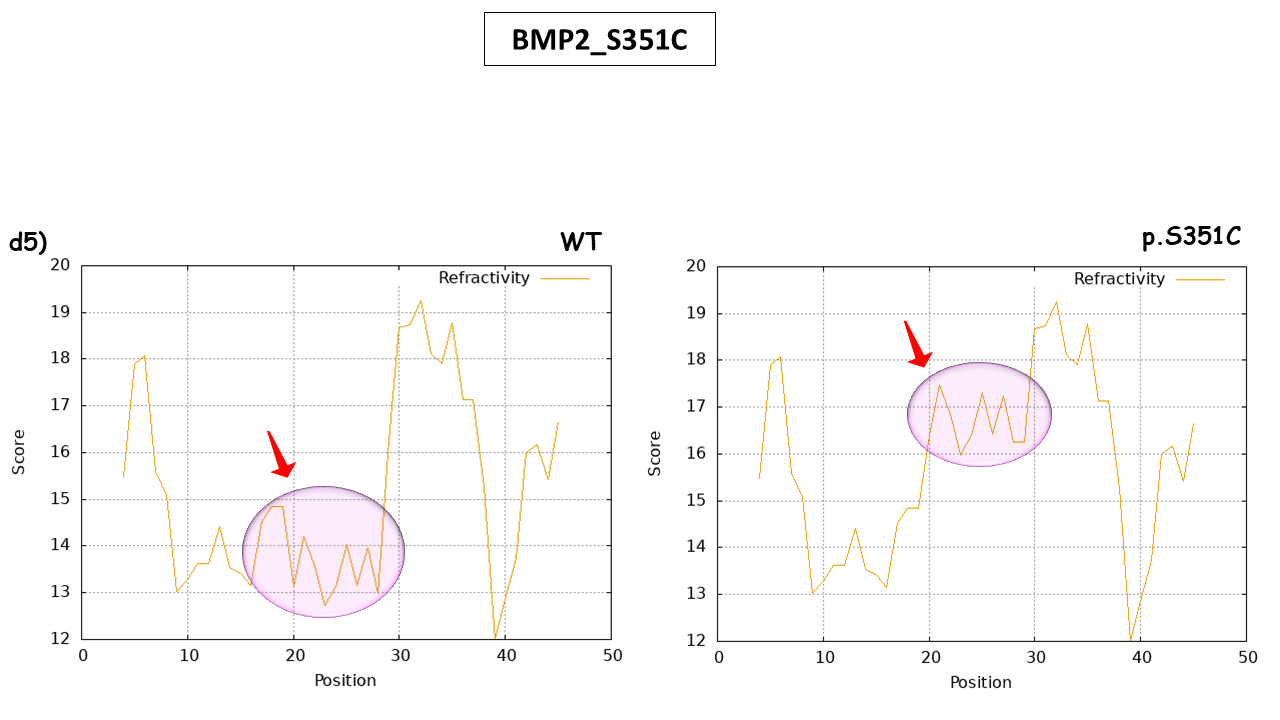

Supplementary Fig. 1. Computational analysis of different physicochemical properties of BMP2 WT vs muteins. (a1-a5) showing tendencies of alpha-helix of WT and all the five muteins, (b1-b5) representing change in ratio hetero end side of muteins over WT, (c1-c5) comparative analysis of relative mutability, (d1-d5) prediction of refractivity. The encircled regions marked with arrow representing the changes. WT= wild-type.

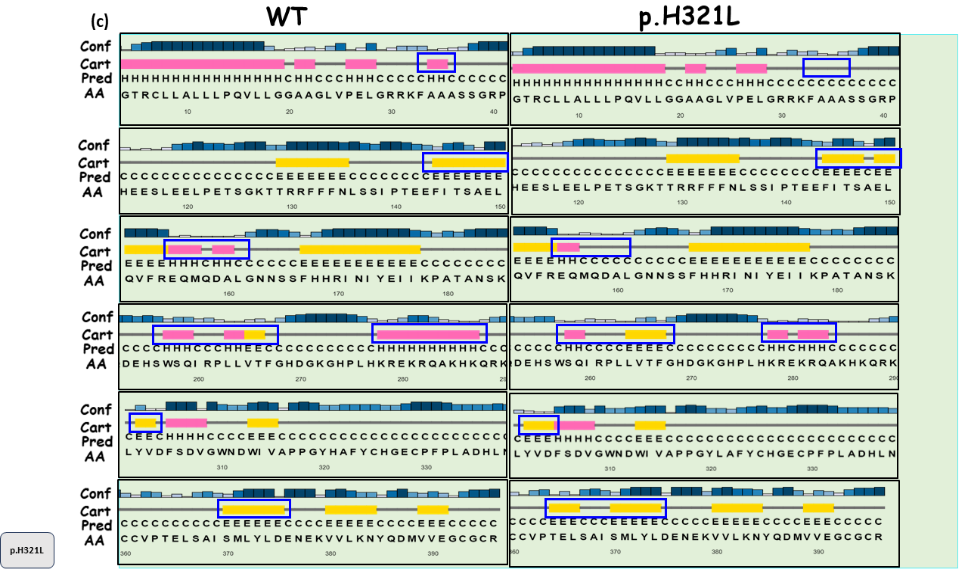

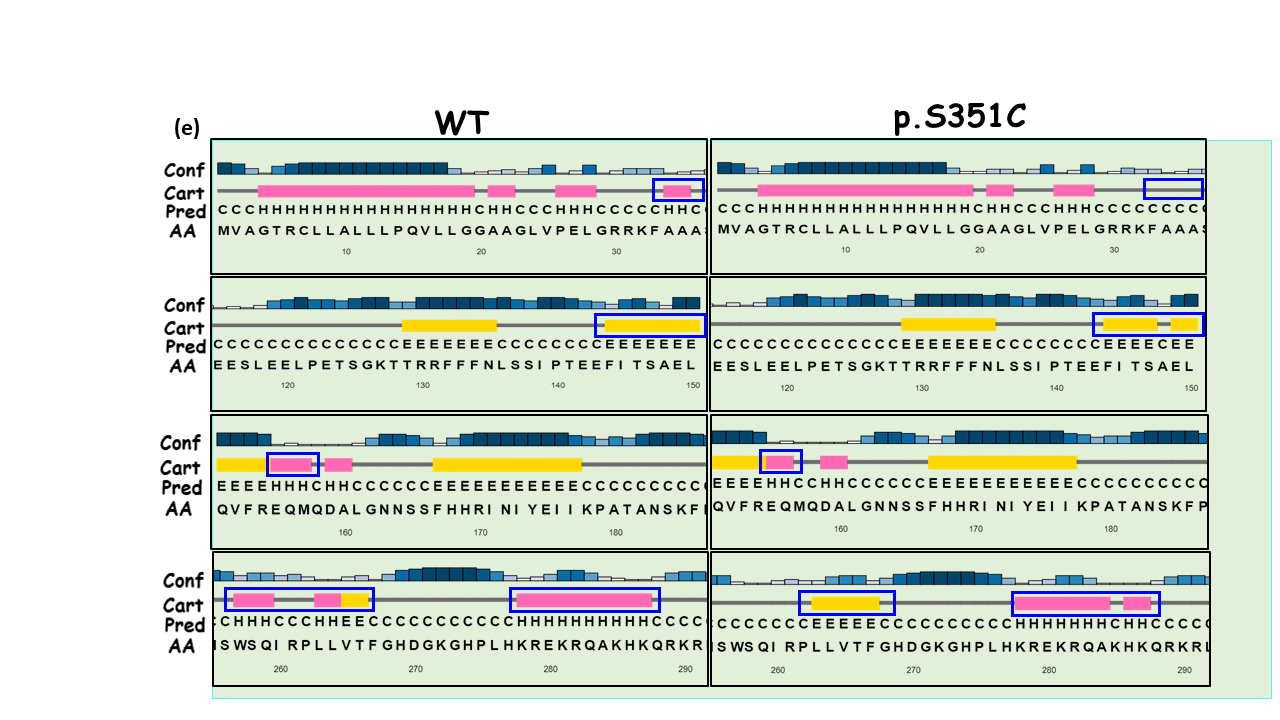

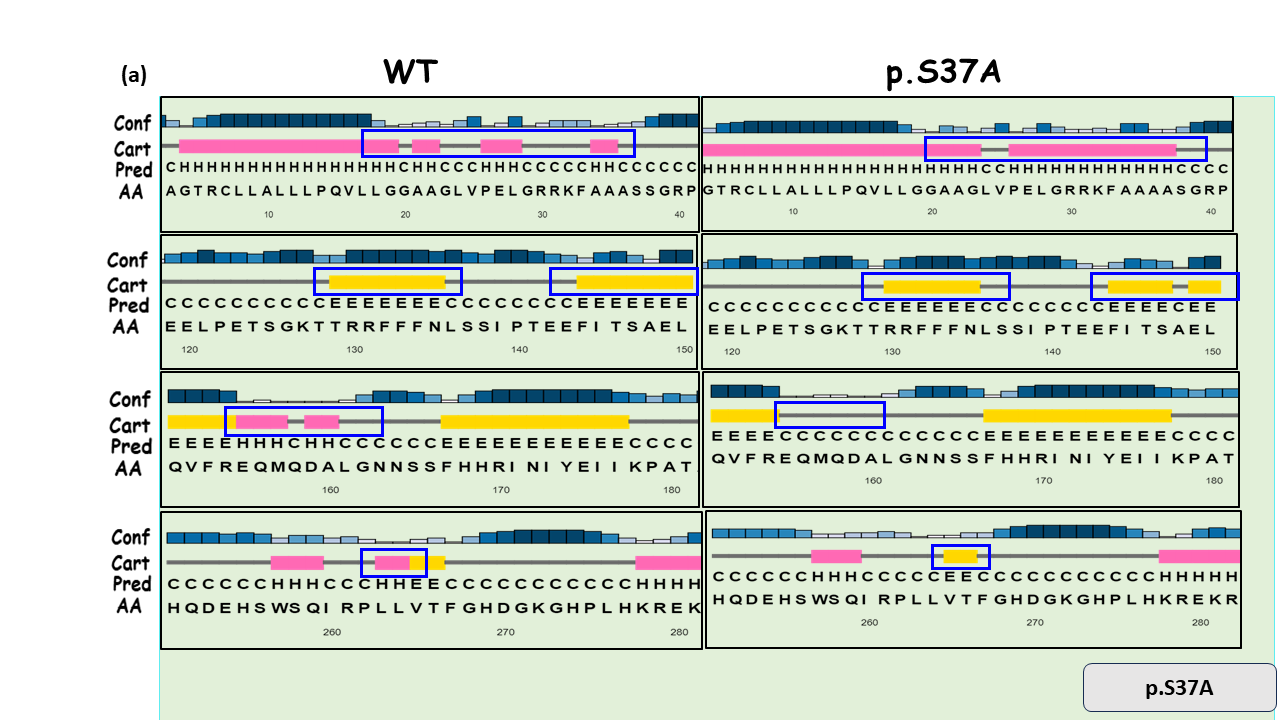

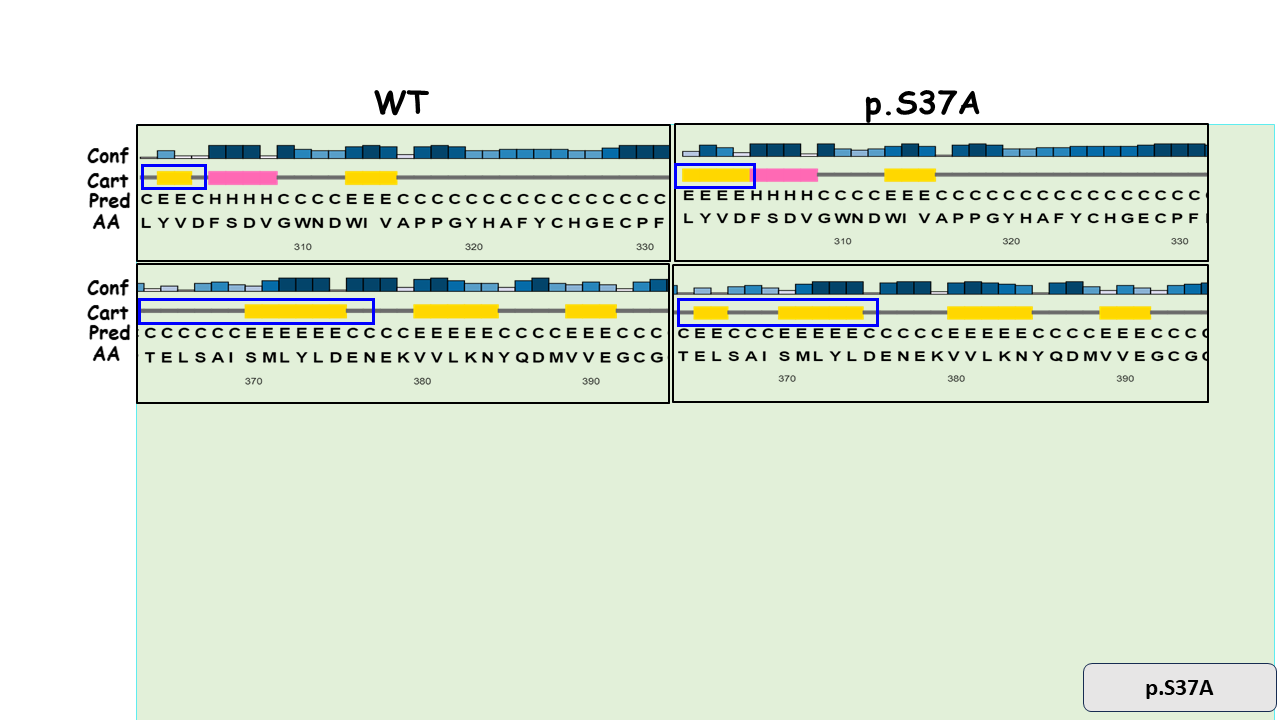

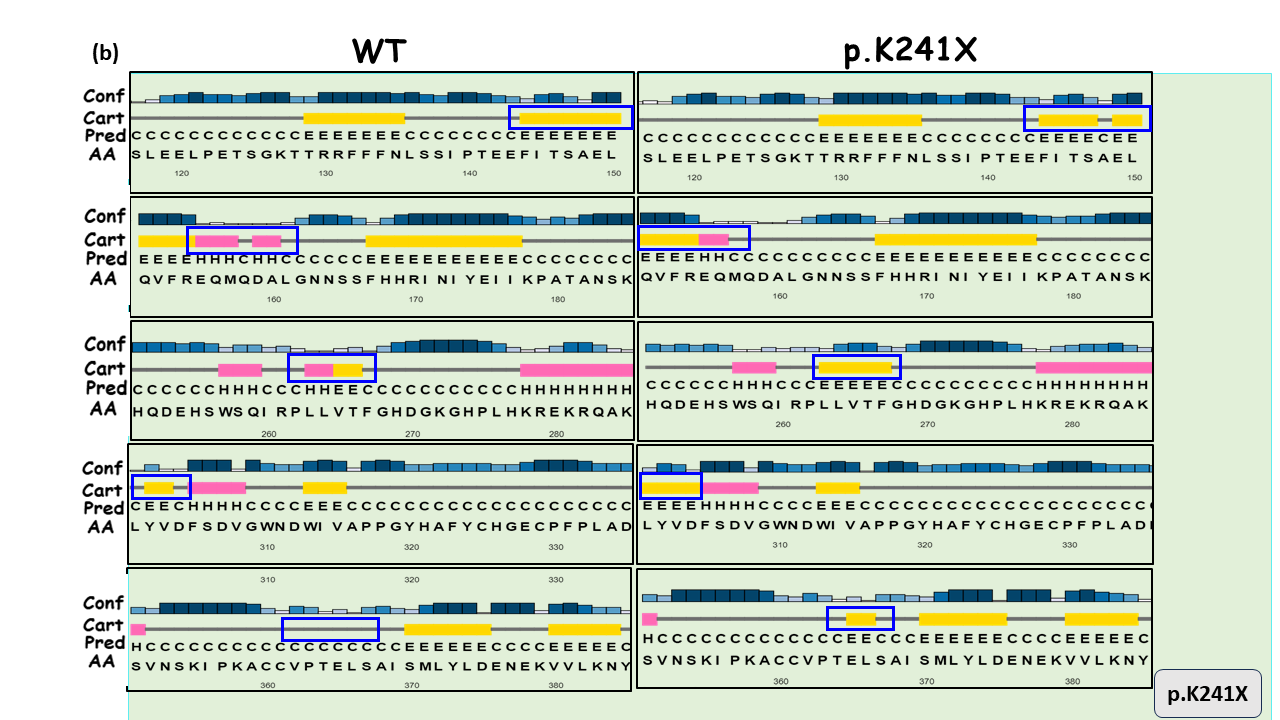

Supplementary Fig. 2. Comparison based on the secondary structures of BMP2 WT and muteins. (a-e) Secondary structures of BMP2 WT and muteins (p.S37A, p.K241X, p.H321L, p.E328K and p.S351C) as predicted by Psipred server. The alterations in structures- helix (pink boxes) and strand (yellow boxes) has been indicated with blue rectangle in both WT and MUTs.

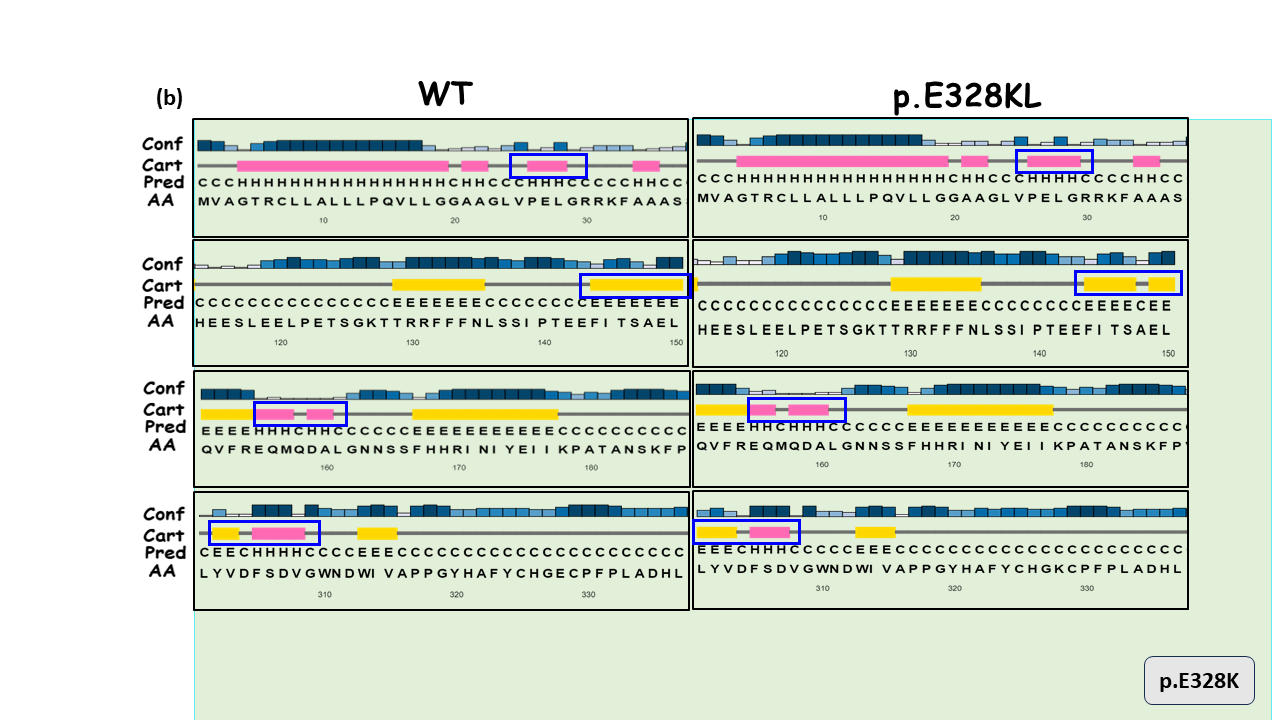

Supplementary Table 1 List of CHD phenotype and its relative frequency in male and females

| **CHD** | **CHD Phenotype** | **Number (%)** |
| --- | --- | --- |
| **Acyanotic defects** | Ventricular septal defects (VSD) | 93 (32.6) |
|  | Atrial septal defects (ASD) | 59 (20.7) |
|  | ASD, VSD | 18 (6.3) |
|  | Patent ductus arteriosus | 7 (2.4) |
|  | Atrio-ventricular canal defect | 6 (2.1) |
|  | Eisenmenger syndrome | 4 (1.4) |
|  | Gerbode defect | 2 (0.7) |
|  | Patent foramen ovale | 1 (0.3) |
| **Cyanotic defects** | Tetralogy/ Pentalogy of Fallot | 44 (15.4) |
|  | Transposition of great arteries | 6 (2.1) |
|  | Dextrocardia | 6 (2.1) |
|  | Tricuspid Atresia | 3 (1.05) |
|  | Ebstein Anomaly | 3 (1.05) |
|  | Double outlet right ventricle | 2 (.07) |
|  | Total/Partial Pulmonary Venous Connection | 2 (.07) |
| **Left/ right obstructive defects** | Bicuspid aortic valve | 12 (4.2) |
|  | Pulmonary stenosis | 6 (2.1) |
|  | Aortic stenosis | 3 (1.05) |
|  | Truncus Arteriosus | 3 (1.05) |
|  | Coarctation of aorta | 1 (0.3) |
| **Others** | Holt Oram Syndrome | 2 (0.7) |
|  | Single Left ventricle | 1 (0.3) |
|  | Double aortic arc | 1 (0.3) |
| **Total** | | **285** |

Supplementary Table 2 Position of changed α-helix and β-sheet caused due to identified variants of *BMP2* as predicted by Psipred server

| Variant | α-helix | β-sheet |
| --- | --- | --- |
| p.Ser37Ala | 20-23^rd^ and 26-36^th^ extension  155-160^th^, 263-264^th^ lost | 130-135^th^; 144-147^th^ shortening  301-304^th^ extension  365-366^th^ lost |
| p.Lys241X | 155-156^th^ shortening  159-160^th^ lost | 148-150^th^ shortening  263-266^th^ extension  301-304^th^ extension  365-366^th^ addition |
| p.His321Leu | 34-35^th^ lost  159-160^th^ lost  264-265^th^ lost  155-156^th^ shortening  198-199^th^ shortening  278-283^rd^ shortening | 144-150^th^ shortening  370-374^th^ shortening |
| p.Glu328Lys | 26-29^th^ extension  155-156^th^ shortening  158-160^th^ extension  305-307^th^ extension | 148^th^ lost  301-303^rd^ extension |
| p.Ser351Cys | 34-35^th^ lost  257-259^th^ lost  155-156^th^ shortening  286-287^th^ shortening | 264-265^th^ addition  148-150^th^ shortening |
